# Global transcription regulation revealed from dynamical correlations in time-resolved single-cell RNA-sequencing

**DOI:** 10.1101/2023.10.24.563709

**Authors:** Dimitris Volteras, Vahid Shahrezaei, Philipp Thomas

## Abstract

Single-cell transcriptomics reveals significant variations in the transcriptional activity across cells. Yet, it remains challenging to identify mechanisms of transcription dynamics from static snapshots. It is thus still unknown what drives global transcription dynamics in single cells. We present a stochastic model of gene expression with cell size- and cell cycle-dependent rates in growing and dividing cells that harnesses temporal dimensions of single-cell RNA-sequencing through metabolic labelling protocols and cell cycle reporters. We develop a parallel and highly scalable Approximate Bayesian Computation method that corrects for technical variation and accurately quantifies absolute burst frequency, burst size and degradation rate along the cell cycle at a transcriptome-wide scale. Using Bayesian model selection, we reveal scaling between transcription rates and cell size and unveil waves of gene regulation across the cell cycle-dependent transcriptome. Our study shows that stochastic modelling of dynamical correlations identifies global mechanisms of transcription regulation.

## 1 Introduction

Transcription occurs in episodic bursts of activity, causing fluctuations in the level of transcripts produced by a single gene [1–3]. The frequency and size of these bursts lead to diverse temporal patterns of gene activity [4, 5], often referred to as cellular noise, that can have a functional role in gene regulatory networks and cell fate decisions [6–14]. The cell cycle has been shown to contribute significantly to transcript variability [15–19], either through random partitioning of molecules at cell division [20–22], modulation of transcription rates [23–27] or through coupling to cell size as a mechanism of transcript homeostasis [28–31]. Yet, it remains elusive what the global drivers of transcription dynamics are on a transcriptome-wide scale.

Single-cell RNA-sequencing (scRNA-seq) has become the de facto standard in profiling transcription patterns. The method allows quantifying distributions of transcript levels across cells that can be harnessed to learn about gene regulation. A few studies have started to investigate how transcriptional processes are regulated through synergising stochastic modelling and inference [32–36]. To this end, stochastic models of reaction kinetics inside living cells are simulated using the Gillespie algorithm or solved analytically and matched with experimental mRNA counts using statistical inference [25, 37–41]. However, a challenge is that not one model fits it all at a genome-wide scale, and identifying transcriptional patterns requires weighing competing model hypotheses against each other that need to be tested against experiments.

In principle, this aim can be achieved using Bayesian inference by computing the posterior distribution over models and their corresponding parameters. The posterior model probability weighs the model likelihood by its prior distribution. Several problems with this approach appear in practice: (i) the likelihood of mechanistic models is intractable or is computationally expensive to optimise [38], (ii) scRNA-seq data is heavily corrupted by technical noise including dropout [42], (iii) absolute kinetic rates are experimentally inaccessible from static snapshots [43].

In the literature, these problems are typically dealt with separately. The first issue is addressed by replacing likelihoods with moment- or simulation-based methods [33–36, 44]. As an example, the authors of [34] present an Approximate Bayesian Computation framework to infer transcriptional burst kinetics from scRNA-seq data. Yet, the above methods either neglect technical noise of sequencing protocols or remain computationally challenging as they need to be performed for several competing models and many genes. The second issue has been addressed by rescaling transcript counts [35, 36, 42, 45], but Bayesian normalisation has been shown to capture distributions more truthfully [46, 47]. To this end, a recent study [36] demonstrated that modelling technical noise can improve inference of transcriptional burst kinetics, yet it does not explore multiple modelling hypotheses. The third issue remains a major obstacle for single-cell analysis, as traditional sequencing protocols can only capture a static snapshot of cells in time, thus prohibiting absolute quantification of transcription and mRNA turnover rates.

Recently, metabolic RNA labelling has become available to temporally resolve conventional single-cell transcriptomics [48]. These protocols allow monitoring of mRNA synthesis and degradation in single cells, thus capturing temporal transcription dynamics [26, 27, 49–54]. This provides new possibilities for modelling as it enables absolute rate quantification, as demonstrated recently for the inference of transcriptional burst kinetics through maximum likelihood methods [55] or MCMC sampling from the posterior distribution [56]. These approaches have either neglected technical noise or have been limited to relatively simple models and have not explored the landscape of the model posterior. It thus remains elusive which models describe gene expression on a transcriptome-wide scale.

Here, we develop a scalable approach to Bayesian inference, normalisation and model selection of bursty gene expression dynamics along the cell cycle. A stochastic model enables absolute quantification of transcription dynamics along the cell cycle, resolving mRNA synthesis, bursting and degradation processes from scRNA-seq data with metabolic labelling and cell cycle reporters [26]. We demonstrate that correlations between unlabelled and labelled mRNA levels can delineate different modes of transcriptional regulation and quantify the contributions from the cell cycle and mRNA synthesis, bursting, and degradation kinetics to the overall cell-to-cell variation. The framework allows us to discover which of these processes are modulated by the cell cycle and identify genes with distinct transcription regulation. Our study thus reveals, for the first time, how genome-wide single-cell transcriptional dynamics is orchestrated with global factors such as cell size and cell cycle.

## 2 Results

### Stochastic modelling captures distinct patterns of bursty transcription dynamics along the cell cycle

We develop a stochastic model to describe the dependence of bursty transcription dynamics on the cell cycle as well as the effects of RNA labelling, according to the scEU-seq protocol. The protocol labels transcripts with a pulse of a certain duration followed by a chase period after which transcripts are sequenced (Figure 1 (A)). We distinguish *pulse* conditions with varying pulse labelling duration and zero chase period, and *chase* conditions with fixed pulse duration (22 hours) and varying chase duration (Supplementary Methods 1.1.2). We assume reaction kinetics based on a two-stage telegraph model of transcription, capturing random gene promoter switching between active and inactive states, followed by synthesis and degradation of labelled and unlabelled mRNA (Supplementary Methods 1.2.1). The rates driving these reactions are assumed to be cell cycle-dependent, and we additionally assume a linear scaling of mRNA synthesis rate with cell size (Figure 1 (B), Methods 1.2.2) [28, 57, 58]. Our model postulates labelling occurs on nascent mRNA [59] and that labelled and unlabelled mRNA are produced with rates proportional to a gene-specific labelling efficiency. We formulate moment equations that describe the temporal dependence of mean and covariance of labelled and unlabelled mRNA on cell cycle progression and cell division dynamics as well as on pulse-chase labelling conditions. These equations track the mRNA statistics over time along a single-cell lineage, following consecutive cell divisions until convergence to a steady state, capturing the random partitioning of molecules at cell division as well as the effect of pulse-chase labelling (Figure 1 (C), Supplementary Methods 1.2.3, 1.2.4, 1.2.6). The cell cycle-dependent changes of mRNA statistics and kinetic rates are described in five distinct cell cycle phases (Supplementary Figure S1, Supplementary Methods 1.1.3).

**Figure 1:**
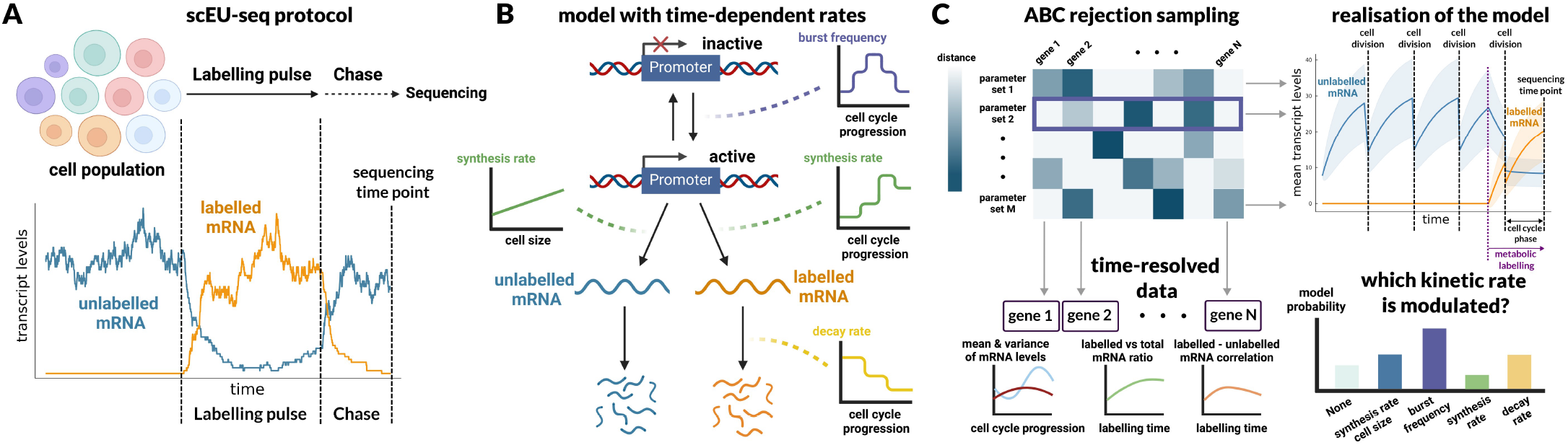
Overview of modelling and inference framework. (A) Illustration of single-cell EU RNA-sequencing (scEU-seq) protocol, developed by [26]. A heterogeneous cell population is pulse-chase labelled prior to sequencing. An example of stochastic transcription of a gene at the single-cell level is shown below, where labelled mRNA is produced and degraded following the timing of pulse-chase labelling. The pulse and chase experiments with multiple durations of the scEU-seq protocol are informative of the timescales of mRNA turnover. (B) Illustration of the general reaction kinetics model describing bursty transcription of unlabelled and labelled mRNA with cell cycle-dependent kinetic rates. The promoter activation rate (burst frequency), synthesis rate and decay rate are cell cycle-dependent. (C) Overview of the ABC inference scheme: Each sampled prior parameter set yields a model realisation which is accepted/rejected based on a comparison with summary statistics for all genes. This provides an estimate of the posterior probability across models.

Our modelling framework makes it possible to test multiple hypotheses on the cell cycle-dependence of kinetic rate parameters and identify genes characterised by distinct underlying kinetics. To this end, we develop an Approximate Bayesian Computation framework [60] to infer the mode of regulation that best describes each gene (Figure 1 (C)), as we explain in more detail in the next section. We define five distinct model classes representing hypotheses about specific cell cycle-dependent regulation of kinetic rates (Figure 2 (A)). Two of the model classes, which we refer to as *constant* models, assume that the synthesis rate either scales linearly with cell size or stays constant, while the rest of the kinetic rates are assumed to be constant along the cell cycle. The other three model classes, which we refer to as *non-constant* models, assume scaling of the synthesis rate with cell size and additionally that one of the kinetic rates, either burst frequency or synthesis rate or decay rate, varies with cell cycle progression. We refer to the three non-constant models as the *burst frequency* model, the *burst size* model and the *decay rate* model, respectively. The two constant models identify genes for which the kinetics are conserved through the cell cycle and distinguish between size-scaling and size-independent expression. The three non-constant models identify genes for which there is a kinetic rate being dominantly modulated through the cell cycle.

**Figure 2:**
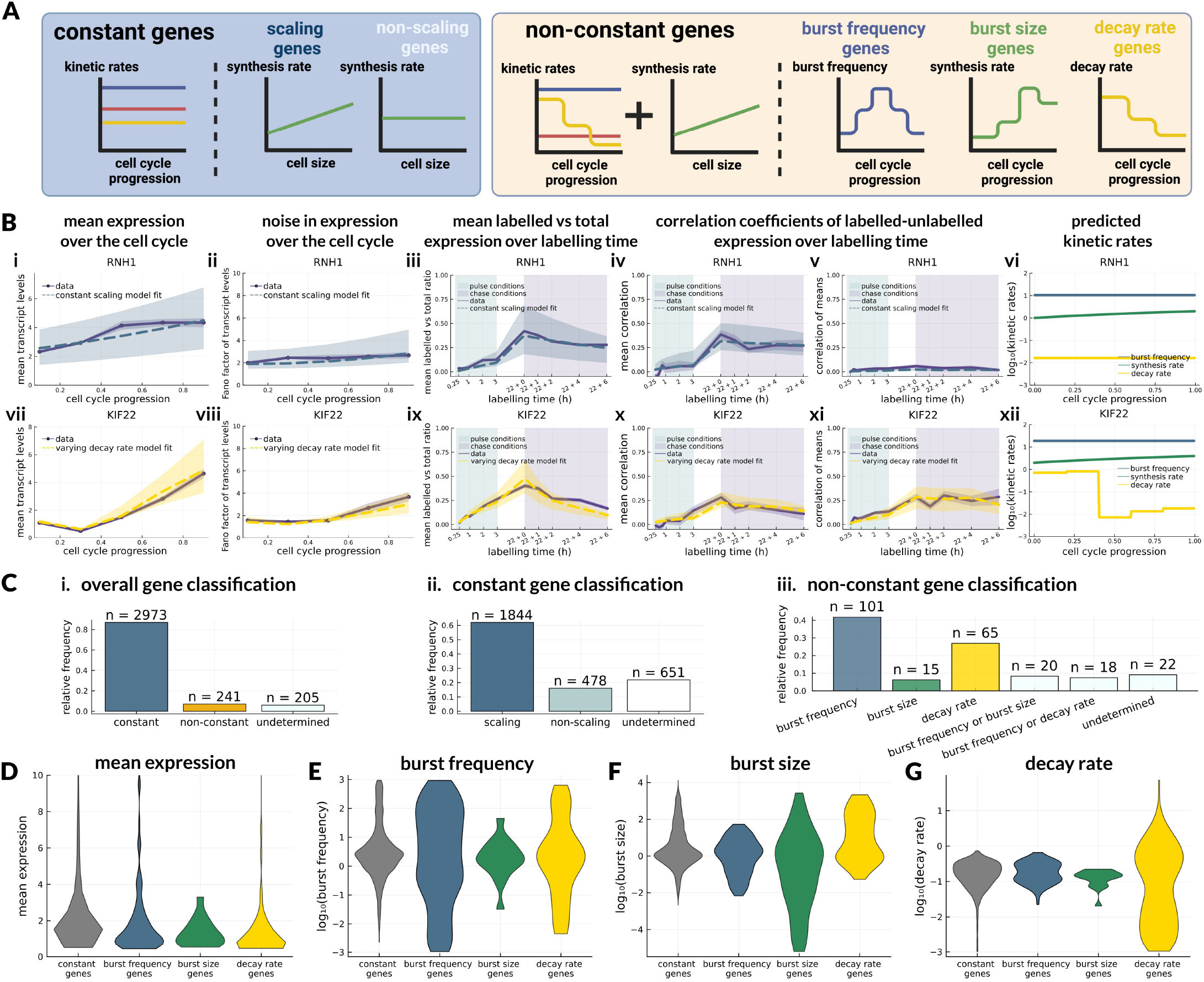
Gene classification with respect to models of kinetics regulation. (A) Cartoon overview of the different model - gene classes. Constant models (blue box) assume that synthesis rate either scales with cell size (scaling genes) or stays constant (non-scaling genes), while all other kinetic rates stay constant along the cell cycle. Non-constant models (yellow box) assume that synthesis rate scales with cell size and additionally one kinetic rate varies with the cell cycle, while the rest of kinetic rates stay constant. (B) Summary statistics used for comparing models and data for an example constant gene (top row) and an example non-constant gene (bottom row), together with estimated kinetic rates for each gene. (C) Overall statistics of gene classification obtained from the ABC model selection. (D-G) Comparison of mean expression and kinetic rate magnitudes across different classes of genes. For each model class (constant and non-constant), proportions of specific sub-classes are shown. For constant, varying burst frequency, varying burst size and varying decay rate genes, we show distributions of (D) mean expression, (E) burst frequency, (F) burst size and (G) decay rate.

We find that these five models yield diverse summary statistics (defined in Tables 1 (A) and 2) and can identify distinct dynamic features of transcription. Example model realisations (Supplementary Figure S2) suggest that noise statistics have a crucial role in model identifiability. For instance, even though the varying burst frequency and burst size models can give rise to overall similar summary statistics, the cell cycle-dependent noise statistics can be used to distinguish between the two models. We also find that the varying decay rate model yields very different patterns compared to the other two non-constant models, suggesting that our model can distinguish regulatory mechanisms of transcription from mechanisms regulating mRNA degradation. Swapping linear for exponential growth did not affect our results as exponential and linear growth are nearly indistinguishable over one cell size doubling (Supplementary Figure S1). Additionally, we observe that correlation coefficients have an essential role in distinguishing constant versus cell cycle-modulated kinetics.

**Table 1:** Definitions of summary statistics. (A) Theoretical model-based definitions. Dependence on the cell cycle is denoted by *τ* and dependence on labelling condition is denoted by *t*. Unlabelled, labelled and total mRNA levels of a single gene are denoted by *u, l* and *x* = *u* + *l*, respectively. (B) Data-based definitions. Notation is explained in paragraph 1.1.5.

|  | A. Model | B. Data |
| --- | --- | --- |
| (1) | $\mathbb{E}(x \mid \tau)$ | $\bar{x}_j(\theta, c) = \frac{1}{N_{\theta, c}} \sum_{i \in \theta \cap c} x_{ij}$ |
| (2) | $\frac{\text{Var}(x \mid \tau)}{\mathbb{E}(x \mid \tau)}$ | $\frac{\frac{1}{N_{\theta, c}-1} \sum_{i \in \theta \cap c} (x_{ij} - \bar{x}_j(\theta, c))^2}{\bar{x}_j(\theta, c)}$ |
| (3) | $\frac{\mathbb{E}_\tau(\mathbb{E}(l \mid \tau, t))}{\mathbb{E}_\tau(\mathbb{E}(u \mid \tau, t) + \mathbb{E}(l \mid \tau, t))}$ | $\frac{\frac{1}{N_t} \sum_{i \in t} l_{ij}}{\frac{1}{N_t} \sum_{i \in t} x_{ij}}$ |
| (4) | $\frac{\mathbb{E}_\tau(\text{Cov}(u, l \mid \tau, t))}{\sqrt{\text{Var}(u \mid t) \text{Var}(l \mid t)}}$ | $\frac{\sum_\theta \frac{N_{\theta, t}}{N_t} \sum_{i \in \theta \cap t} (u_{ij} - \bar{u}_j(\theta, t))(l_{ij} - \bar{l}_j(\theta, t))}{\left( \frac{1}{N_t} \sum_{i \in t} (u_{ij} - \bar{u}_j(t))^2 \frac{1}{N_t} \sum_{i \in t} (l_{ij} - \bar{l}_j(t))^2 \right)^{1/2}}$ |
| (5) | $\frac{\text{Cov}_\tau(\mathbb{E}(u \mid \tau, t), \mathbb{E}(l \mid \tau, t))}{\sqrt{\text{Var}(u \mid t) \text{Var}(l \mid t)}}$ | $\frac{\sum_\theta \frac{N_{\theta, t}}{N_t} \left( \bar{u}_j(\theta, t) - \sum_\theta \frac{N_{\theta, t}}{N_t} \bar{u}_j(\theta, t) \right) \left( \bar{l}_j(\theta, t) - \sum_\theta \frac{N_{\theta, t}}{N_t} \bar{l}_j(\theta, t) \right)}{\left( \frac{1}{N_t} \sum_{i \in t} (u_{ij} - \bar{u}_j(t))^2 \frac{1}{N_t} \sum_{i \in t} (l_{ij} - \bar{l}_j(t))^2 \right)^{1/2}}$ |

**Table 2:** Interpretation of summary statistics.

|  | Interpretation |
| --- | --- |
| (1) | cell cycle-dependent mean of mRNA levels |
| (2) | cell cycle-dependent Fano factor of mRNA levels |
| (3) | ratio of mean labelled vs total mRNA levels over labelling time |
| (4) | cell cycle-averaged correlation of labelled-unlabelled mRNA levels over labelling time |
| (5) | correlation of cell cycle-dependent labelled & unlabelled means over labelling time |

### Approximate Bayesian inference reveals global patterns of transcription kinetics regulation

Motivated by the ability of the model to produce distinct summary statistics between different underlying modes of kinetics regulation, we develop an inference framework to infer these modes of regulation along with gene-specific kinetic parameters from scEU-seq data. A main challenge here is that inference requires optimising over a large parameter space across all five modelling hypotheses for each gene in the data separately. Therefore, using a maximum likelihood, optimisation-based or Markov Chain Monte Carlo approach would be computationally inefficient. For that reason, we develop a simulation-based inference framework based on ABC rejection sampling, which allows to test multiple modelling hypotheses and millions of parameter sets against the entire set of genes in a highly parallel manner (Figure 1 (C)). Given a parameter set sampled from a prior distribution and a realisation of the model, we generate a collection of summary statistics (see Tables 1, 2 and Supplementary Methods 1.4) and simultaneously compare them to summary statistics of all genes in the data (Supplementary Methods 1.5.1). Additionally, using an Approximate Bayesian framework allows to perform model selection across several models with different levels of complexity (number of parameters) [61], *i*.*e*. to assign probabilities to the various hypotheses, and decide which kinetic model is most likely to describe each gene. A crucial challenge for inference is that scEU-seq data are characterised by low mRNA capture efficiency [26], meaning that summary statistics of the data and the model are not directly comparable. We address this issue by modelling cell-specific capture efficiencies and downsampling the moments to enable reliable inference [36, 47, 62], as we describe in detail in the next sections.

In Figure 2 (B) we show example posterior fits over the different summary statistics (see Tables 1 and 2) for a constant and a non-constant model. We observe that the mean expression of the gene *RNH1* shows a monotonous, approximately linear increase, while its Fano Factor stays relatively constant over the cell cycle (Figures 2 (B i-ii)). Even though the constant model shows a linear increase in both mean expression and Fano factor, the posterior model fit agrees well with the data. The ratio of mean labelled vs total expression of *RNH1* increases with pulse labelling time and decreases along chase conditions as the chase duration increases (Figure 2 (B iii)). The model behaves similarly and we see that the data lies well within the posterior model fit. The mean (cell cycle-averaged) correlation of labelled-unlabelled expression follows a similar pattern with the ratios and even though the data exhibits a more noisy behaviour, its error bars lie within the posterior model fit (Figure 2 (B iv)). The correlation of cell cycle-dependent means is almost zero across labelling conditions for both model and data (Figure 2 (B v)).

The gene *KIF22* shows a large variation in mean and Fano factor of expression over the cell cycle (Figures 2 (B vii-viii)) as we see a strong increase towards the late cell cycle stages. The non-constant decay rate model explains these patterns well as the error bars of the data lie within the posterior fit. The ratio of mean labelled vs total expression and the mean correlation of labelled-unlabelled expression of *KIF22* show similar behaviour with the respective statistics of *RNH1*, and even though they appear more noisy than the model, we obtain a good overall posterior fit (Figures 2 (B ix-x)). The correlation of means is significantly higher compared to the *RNH1* case, and even though it is also very noisy, it is captured well by the non-constant posterior fit (Figure 2 (B xi)). These model fits show that *RNH1* is explained by constant kinetics along the cell cycle and its synthesis rate scales linearly with cell size (Figure 2 (B vi)). On the other hand, the gene *KIF22* is explained by non-constant kinetics as its decay rate varies with cell cycle progression (Figure 2 (B xii)). Overall, our inference framework provides us with approximate posterior distributions of gene-specific kinetic parameters (Supplementary Figure S3). From this example we see that synthesis and decay rate are better identified compared to burst size and burst frequency. Interestingly, we observe that burst size and frequency are negatively correlated as well as that decay rate estimates are independent from the rest of the parameters.

After performing parameter inference over all 3419 genes that we selected for the analysis (Supplementary Methods 1.1.4), we find that the majority (87%) are explained by constant models, and about 7% of genes are assigned to non-constant models (Figure 2 (C i)). Within the subset of genes that are assigned to constant models (*constant* genes), we find that the majority (62%) are assigned to the size-scaling synthesis rate model (*scaling* genes) and a smaller fraction of genes (16%) are assigned to the non-scaling synthesis rate model (*non-scaling* genes), while for the rest of constant genes we get an uncertain model choice outcome (Figure 2 (C ii)). Among the subset of genes assigned to non-constant models (*non-constant* genes), we find two major groups assigned to the burst frequency model (41%) and the decay rate model (27%), respectively, while a small fraction of genes are assigned to the burst size model (Figure 2 (C iii)). For the rest of non-constant genes, we find two noteworthy groups of genes, one that is described well by both the burst frequency and the burst size model and one that is described well by both the burst frequency and the decay rate model. For the remaining genes (6%) we either get an uncertain model selection outcome or all the models performed poorly.

Furthermore, our inference reveals general patterns of mRNA kinetics across the different groups of genes. We observe that genes in all groups span a wide range of mean expression levels, while we observe that constant genes have a slightly higher median expression level compared to non-constant genes (Figure 2 (D)). We find that burst frequency is highly variable across all constant and non-constant genes, while burst frequency genes exhibit the strongest variation (Figure 2 (E)). Additionally, cell cycle-regulated burst size shows the largest variation, reaching very low levels compared to genes with constant burst size (Figure 2 (F)). We similarly see large variation in decay rate among decay rate genes, while for the rest of the genes the decay rate is constrained in similar levels (Figure 2 (G)).

We also find that cell cycle-dependent regulation of decay rate impacts mRNA half-life. The majority of genes have median half-lives of approximately 4-5 hours, while the median mRNA half-life of genes with cell cycle-dependent decay rate is longer than 10 hours (Supplementary Figure S4 (A) and (D)). The predicted half-lives are thus shorter than the cell cycle duration (Supplementary Methods 1.2.6). After comparing our estimates of mRNA half-life with two other metabolic labelling studies (sci-fate [52] and scSLAM-seq [49]), we find that the estimated half-lives of constant genes agree well with the sci-fate predictions (Spearman’s *ρ* ≈ 0.9, Supplementary Figure S4 (B)), however a lower correlation is observed in the case of non-constant genes (Supplementary Figure S4 (E)). We find good agreement between our constant gene half-lives and the scSLAM-seq predictions (Spearman’s *ρ* ≈ 0.7, Supplementary Figure S4 (C)), while there is a poor agreement in the case of non-constant genes (Supplementary Figure S4 (F)). These discrepancies in the case of non-constant genes indicate that modelling cell cycle-dependence improves the estimation of mRNA half-life. The overall discrepancies could in addition be attributed to differences between the scEU-seq, sci-fate and scSLAM-seq metabolic labelling protocols [48] or to the fact that these estimations are based on different cell lines.

We validated the correctness and robustness of our inference and model selection framework. We found that doubling the number of simulations does not change our conclusions on model classification (Supplementary Figure S5 (A)). We also found that when including more complex non-constant models (where more than one parameter varies with the cell cycle), our inference still robustly selects constant versus non-constant models, but there is uncertainty in distinguishing between non-constant models (Supplementary Figure S5 (B)). This suggests that richer data are necessary to identify more complex mechanisms. Moreover, we show that our model accurately quantifies 3rd order moments of gene expression, reflecting the predictive capacity of our model (Supplementary Figure S6 (A-B)), which explains both biological and technical noise contributions (Supplementary Figure S6 (C-D)). Thus, including higher-order moments would not improve inference accuracy further. We tested the independence of our summary statistics showing negligible pairwise correlations (Supplementary Figure S7). Finally, we performed inference on synthetic data, generated using Gillespie simulations (Supplementary Methods 1.6), and found that the model accurately recovers models and their parameters (Supplementary Figure S8). In summary, these results suggest that our choice of summary statistics achieves good inference accuracy and model selection.

### Modelling capture and labelling efficiency allows to uncouple technical and biological noise

The technical biases associated with sequencing protocols make the task of modelling scRNA-seq data challenging. Building a stochastic model that accurately explains the observed variability in the data, requires that the various biological and technical contributions to the total noise are carefully accounted for. We here demonstrate that the relationship between unlabelled and labelled mRNA levels of a single gene can reveal how different noise components contribute to the total transcript variability. In the absence of technical noise, the total variance in transcript levels *x* = *u* + *l* of a gene (*u*: unlabelled, *l*: labelled) in a time-resolved single-cell snapshot can be partitioned as follows (see details in Supplementary Methods 1.7):

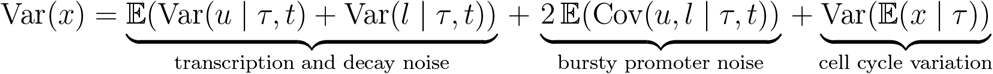

where *τ* denotes the cell cycle phase and *t* the pulse-chase labelling time (condition). Averaging the cell cycle- and labelling time-dependent transcript variances gives a measure of the noise due to mRNA synthesis and degradation kinetics (1st term). This term includes variation due to both active degradation and dilution, where the latter stems from stochastic partitioning at cell division. The upstream noise due to bursty gene promoter can be measured by averaging the cell cycle- and labelling time-dependent covariances of unlabelled and labelled transcripts (2nd term). The extrinsic noise due to cell cycle variation can be measured by the variance of the cell cycle-dependent mean transcript levels (3rd term). The bursty promoter noise and the cell cycle variation contribute to correlating the unlabelled-labelled transcript levels, while transcription and degradation kinetics noise reduces these correlations (Figure 3 (A)). This decomposition is challenging to quantify directly from the data as each component would be corrupted by multiplicative technical noise effects distorting their relative proportions (Supplementary Methods 1.7.1).

**Figure 3:**
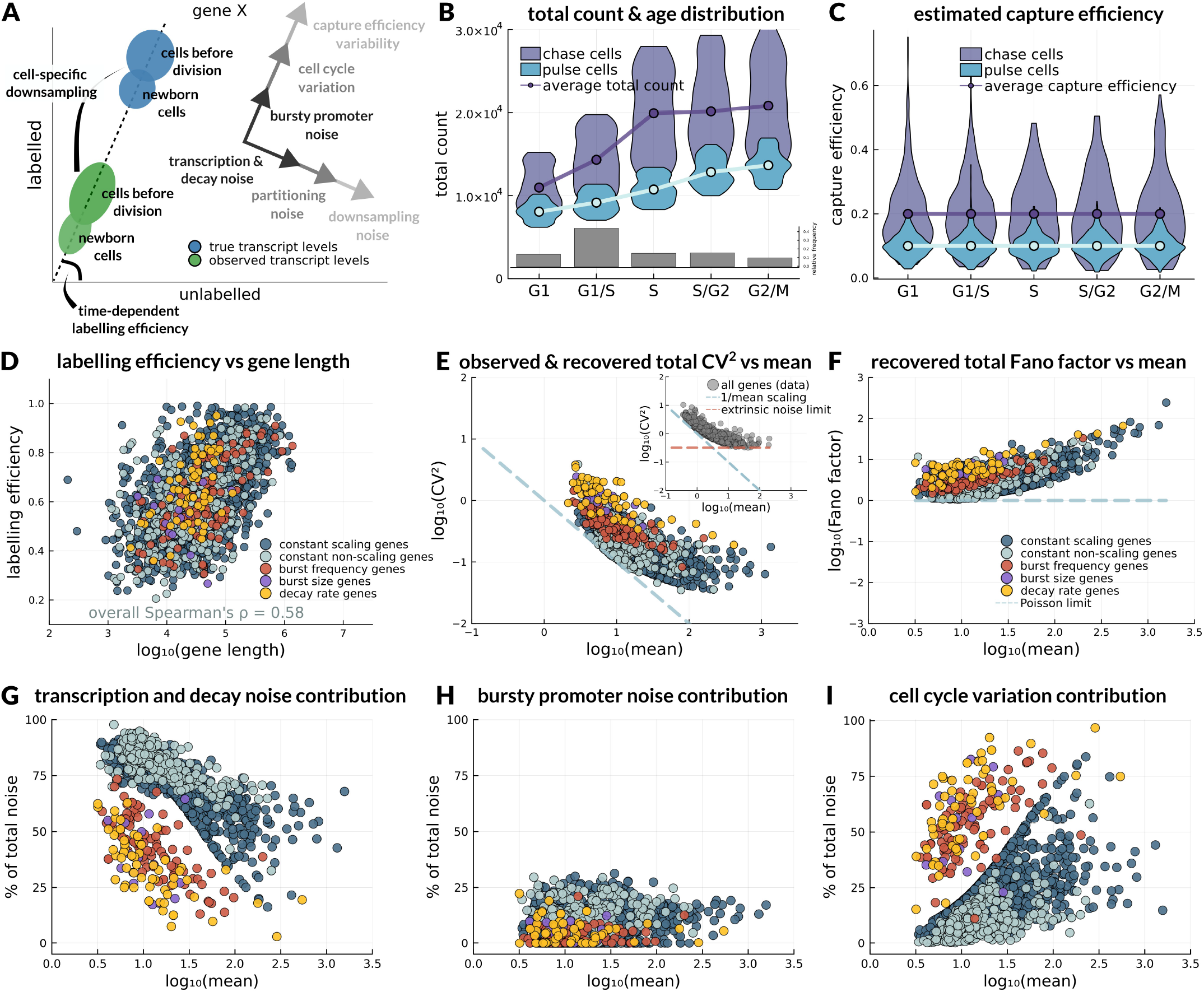
Biological and technical effects on transcriptional noise and biological noise decomposition. (A) Cartoon scatter plot of unlabelled-labelled mRNA, where the observed transcript levels (green ellipses) is a noisy version of the true transcript levels (blue ellipses) due to cell-specific downsampling. The vectors describe different noise components that contribute to the total transcript variability by correlating or decorrelating the unlabelled-labelled mRNA levels, illustrated by lengthening or widening the ellipses, respectively. Bursty promoter noise, cell cycle variation (biological) and capture efficiency variability (technical) contribute to correlating unlabelled and labelled transcript levels, while transcription & mRNA degradation noise, partitioning noise due to cell division (biological), and cell-specific downsampling (technical) contribute to decorrelating transcript levels. (B - C) Uncoupling biological and technical effects on transcript variability: (B) Total mRNA count distributions per cell cycle phase and age distribution contribute to biological variability. The average total count is shown to approximately double over the course of the cell cycle. Calculation of the empirical age distribution is described in Supplementary Methods 1.1.3. (C) Estimated capture efficiency distributions per cell cycle phase. The distributions are shown to be independent of cell cycle progression. (D) Dependence of inferred gene-specific labelling efficiency on gene length. (E-F) Observed and recovered noise-mean relationships: (E) shows the total CV_2_ as observed in the data (inset scatter plot in grey) and as recovered by the different models. (F) shows the recovered total Fano factor as given by the model after removing technical noise. (G-I) Model-based biological noise decomposition: proportional contributions of the different biological noise sources to the total noise (Supplementary Methods 1.7).

We therefore devise an approach to account for technical noise in our modelling framework. This allows us to construct reliable summary statistics that are directly comparable with the noisy data. Quantifying what part of transcript variation comes from technical noise requires access to the capture efficiency of the sequencing experiment, however this information is not provided by the scEU-seq protocol. In principle, actual and observed mRNA counts can be connected via a binomial downsampling model that depends on a cell-specific capture efficiency, but in our case, the dependence on cell size and cell cycle needs to be considered (Figure 3 (B), Supplementary Methods 1.3). We estimate the capture efficiencies directly from the data. The idea is that the total mRNA count of a cell is proportional to cell size and it increases along the cell cycle so it provides an estimate of the relative capture efficiency of the cell within its cell cycle phase. Also, this estimate differs between pulse and chase conditions (Supplementary Methods 1.3.1). We show that capture efficiency distributions per cell cycle phase do not scale with cell cycle progression (Figure 3 (C)). Our approach hence decouples cell size and age variability from technical noise effects on total mRNA count. Moreover, it normalises the total count and sample size differences between pulse and chase conditions (Supplementary Methods 1.3.1). Having obtained the capture efficiency estimates, we can correct the model outputs by computing downsampled versions of moments (Supplementary Methods 1.3.2).

An additional technical noise source at the single-gene level is the labelling efficiency of the pulse-chase experiments, as RNA labelling might not impact genes uniformly. To address this, we model labelling efficiency as a gene-specific parameter that is fitted simultaneously with the kinetic rates. We find that the inferred labelling efficiency shows significant variation across genes as well as that it correlates with gene length, demonstrating the importance of estimating it as a gene-specific parameter (Figure 3 (D)). It is also evident that this trend is independent of the selected gene-specific models, thus validating that it does not influence model selection.

### Bayesian inference reconstructs the transcriptional noise landscape

Our approach of uncoupling technical and biological noise makes it possible to estimate the true mean expression levels and the true levels of transcriptional noise. The observed total noise (CV^2^) for all 3419 genes decreases with mean expression but stays above the Poisson limit (Figure 3 (E) inset), and deviates from the inverse mean scaling for highly expressed genes. This deviation at high expression levels reflects extrinsic noise attributed to both biological and technical effects. Informing the model with the point estimates from the inferred kinetics posteriors, we recover and visualise the true noise-mean relationship (Figure 3 (E)). The dependence of noise on the mean recovered by the model follows a similar pattern to the data, but the noise is an order of magnitude lower and deviates less from the inverse mean scaling law as these deviations are purely due to biological noise. We observe distinct noise patterns among groups of genes explained by different models, with cell cycle-dependent genes commonly displaying higher noise levels than constant genes. Furthermore, using the Fano factor to quantify the noise-mean relationship, we observe that the increase in the mean is associated with increasing deviation from the Poisson limit due to extrinsic noise (Figure 3 (F)).

Next, we use the noise decomposition formula to quantify how the different biological noise sources contribute to the total transcript variability (Figure 3 (G-I)). We find that transcription and decay kinetics and cell cycle variation are the most prominent sources of noise. Interestingly, for most constant genes the largest proportion of noise is explained by mRNA synthesis and decay kinetics (Figure 3 (G)) that contribute to more than 50% of the total noise. On the other hand, cell cycle variation is the dominant source of noise for most non-constant genes (Figure 3 (I)). The bursty promoter noise is less prominent across genes, however not negligible as in a considerable number of genes it explains more than 20% of the noise. Constant and non-constant genes can be distinguished by their absolute levels of cell cycle variation (Supplementary Figure S9).

To gain mechanistic insights into the determinants of transcriptional noise, we illustrate how the different noise sources depend on the kinetic rates. We find that the decay rate is a key determinant of all noise components. Transcription and decay noise and bursty promoter noise increase with decay rate (Supplementary Figure S10 (A-B)). The cell cycle variation of constant scaling genes is constant and hence does not depend on the decay rate, however the cell cycle variation of non-scaling genes decreases with decay rate (Supplementary Figure S10 (C)). These observations agree with model predictions reported in [21] and [29]. We find that transcription and decay noise is relatively independent of burst size (Supplementary Figure S11 (A)). Additionally, bursty promoter noise increases with burst size for large decay rates (Supplementary Figure S11 (B)), while it decreases with burst frequency for small decay rates (Supplementary Figure S11 (C)). These findings highlight that quantifying absolute kinetic rates in combination with model-based noise decomposition reveals the sources of transcriptional variation.

### The model identifies scaling properties of gene expression with cell size

We next focus on genes with constant kinetics and we find that the model reveals key differences between size-dependent and size-independent gene expression. As an example, *RAB7A*, a gene that is involved in intracellular vesicle trafficking and growth-factor-mediated signalling, is explained by a constant scaling model and *MGST1*, a gene responsible for protecting endoplasmic membranes from oxidative stress, is explained by a non-scaling model (Figure 4 (A)). *RAB7A* shows an increase in the mean and Fano factor of transcript levels along the cell cycle (Figure 4 (A i-ii)), which is explained by a size-scaling mRNA synthesis rate (Figure 4 (A iii)), while the mean transcript levels and Fano factor of *MGST1* stay relatively constant during cell cycle progression (Figure 4 (A iv-v)), which is explained by a constant synthesis rate (Figure 4 (A vi)).

**Figure 4:**
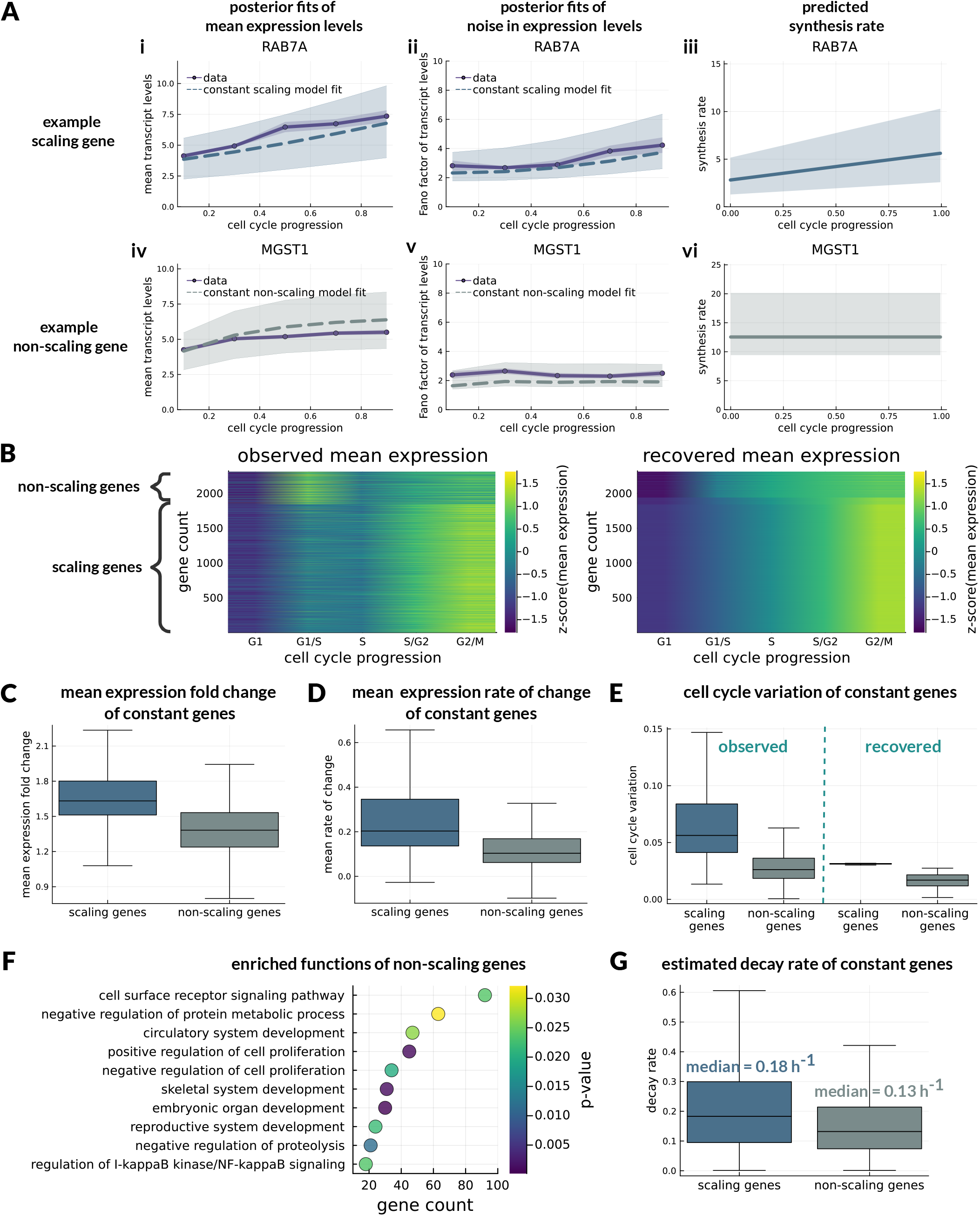
Constant genes and size-scaling properties of gene expression. (A) Posterior model fits for a scaling and a non-scaling gene. Panels (i)-(ii) and (iv)-(v) show cell cycle-dependent mean and Fano factor of transcript levels, and panels (iii) and (vi) show posterior estimates of scaling and non-scaling synthesis rate. (B) Clustered heatmaps of normalised mean expression along the cell cycle for constant genes, as observed in the data (left) and as recovered by the model (right), after removing technical noise effects. (C-E) Differences between scaling and non-scaling genes in cell cycle-dependent expression: (C) shows differences in the observed fold change of mean expression between G1 and G2/M phase, (D) shows differences in mean rates of change of expression along consecutive cell cycle phases and (E) shows differences in variability due to cell cycle-dependence, both observed in the data and recovered by the model. (F) Enriched GO terms of non-scaling genes. (G) Differences in decay rate distributions of scaling and non-scaling genes.

We investigate global size-scaling properties of gene expression. We find that the mean expression of scaling genes increases monotonically during the cell cycle, while the mean expression of non-scaling genes reaches a steady state towards the end of cell cycle progression. This can be observed in clustered heatmaps of both observed and recovered (after technical noise removal) mean expression of all constant genes (Figure 4 (B)). To validate this observation, we quantify changes in mean expression of constant genes along the cell cycle. We find that both the fold change of expression between early and late cell cycle and the mean rate of change of expression along cell cycle progression are higher for scaling compared to non-scaling genes (Figures 4 (C-D)). Additionally, we quantify variation due to cell cycle-dependence, including variation observed in the data and variation recovered from the model after technical noise correction (see Supplementary Methods 1.7). We find that in both cases, cell cycle variation of size-scaling expression is higher compared to non-scaling (Figure 4 (E)). This suggests that the combined signatures of mean expression and noise levels identify the scaling with cell size.

We analysed which gene functions were associated with non-scaling gene expression using the enrichment analysis provided by the DAVID bioinformatics database [63]. We find that the majority of non-scaling genes are involved in cell surface receptor signalling. Other enriched sets include protein metabolic processes, regulation of cell proliferation, tissue and system development and proteolysis. Finally, we asked how scaling of transcription with cell size relates to mRNA degradation. We find that scaling genes have overall higher decay rates compared to non-scaling genes (Figure 4(G)). As a consequence, scaling genes are characterised by shorter mRNA half-life, with their median mRNA half-life being shorter by more than one hour compared to non-scaling genes (Supplementary Figure S4 (A)).

### The model predicts patterns of cell cycle-dependent kinetics regulation

Next, we focus on genes with non-constant kinetics and we find that the model identifies three groups of genes, each characterised by a distinct mode of kinetics regulation during the cell cycle. For these groups of genes, either burst frequency, burst size or degradation rate are primarily modulated over the cell cycle, while the rest of kinetic rates stay constant. Posterior model fits for *CDK4, PCNA* and *CDK1*, which are genes with key functional roles in cell cycle control, display good agreement with the data (Figure 5 (A)). The variations in mean and Fano factor of *CDK4, PCNA* and *CDK1* transcript levels along the cell cycle (Figure 5 (A i-ii, iv-v, vii-vii)) are explained by a particular modulation of their kinetic rates (Figure 5 (A iii, vi, ix)). Interestingly, in all three cases the predicted kinetics show a peak in a particular cell cycle phase. The increase in mean and noise of expression of these genes in the late part of the cell cycle is achieved by a sharp increase in burst frequency or burst size about S phase, or by a sharp decrease in decay rate after G1 phase, respectively.

**Figure 5:**
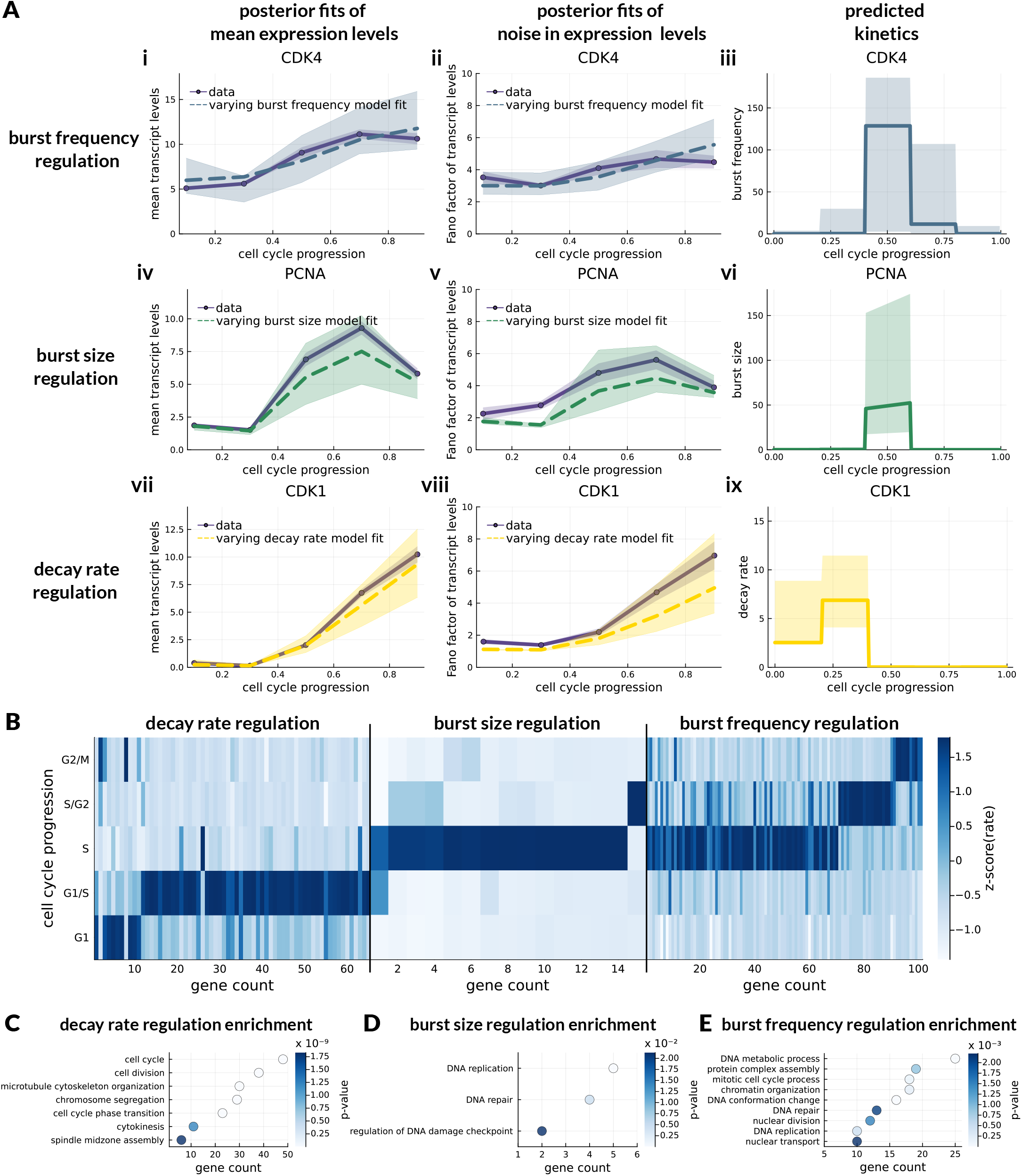
Non-constant genes and cell cycle-dependent regulation of kinetic rates. (A) Posterior model fits for three example non-constant genes. Panels (i)-(ii),(iv)-(v) and (vii)-(viii) show cell cycle-dependent mean and Fano factor of transcript levels, and panels (iii), (vi) and (ix) show posterior estimates of cell cycle-modulated kinetic rates. (B) Clustered heatmap of cell cycle-modulated kinetic rates: the different sections of the heatmap correspond to decay rate genes, burst size genes and burst frequency genes. (C-E) Enriched GO terms associated with non-constant genes: (C) shows GO terms linked with decay rate regulation, (D) shows GO terms linked with burst size regulation and (E) shows GO terms linked with burst frequency regulation.

To obtain comprehensive insights into the global regulation of kinetics during the cell cycle, we perform hierarchical clustering of each set of varying kinetic rates based on cosine similarity (Figure 5 (B)). Strikingly, all emerging clusters are characterised by a unique peak in a particular cell cycle phase. Combining all clusters together, we observe a wave-like pattern in the overall kinetics regulation throughout the cell cycle. We find that varying decay rate shows peaks in the early stages of the cell cycle, while varying burst size and frequency show peaks in S, G2 and M phase. This suggests distinct regulatory strategies of transcription and degradation during the cell cycle. This non-trivial clustering of genes based on their kinetics would not be possible to extract from mean expression dynamics alone, as the different gene groups show similar expression profiles, with an increase towards the late cell cycle stages (Supplementary Figure S12).

We also find, using gene enrichment analysis, that each mode of kinetics regulation is associated with a particular set of functions related to cell cycle control and cell division (Figures 5 (C-E)). Genes with varying decay rate are found to be involved in the regulation of G2/M phase transition and cell division, including chromosome segregation, spindle assembly and cytokinesis (Figure 5 (C)). Genes with varying burst size are primarily involved in S phase regulation and DNA replication, including DNA repair and damage checkpoint control (Figure 5 (D)). Varying burst frequency genes are associated with a variety of cell cycle control functions, including DNA replication, DNA repair and nuclear division (Figure 5 (E)). Interestingly, burst frequency regulation is also associated with other cellular functions such as protein complex assembly, chromatin organisation and DNA conformation changes, suggesting a potential link of these processes with the cell cycle. Overall, these results indicate that biological functions responsible for cell cycle regulation could be achieved by the coordinated activation of kinetic rates in particular cell cycle phases.

## 3 Discussion

We developed a comprehensive modelling framework to reveal global regulatory mechanisms of mRNA transcription and degradation during the cell cycle from time-resolved transcriptomics data. Our stochastic model integrates cell cycle progression and pulse-chase labelling to reveal different modes of dynamic regulation in mRNA control. Modelling capture and labelling efficiency allowed us to deconvolve technical and biological noise of the data and reveal the true sources of cellular variation that stem from distinct mechanisms of transcription regulation.

A central result of our analysis is that patterns of cell cycle- and cell size-dependent regulation of transcription dynamics are distinguished through dynamic correlations of metabolic labels. In particular, we showed that correlations between unlabelled and labelled mRNAs quantify transcriptional bursting, while transcript noise statistics quantify mRNA synthesis and degradation kinetics (Supplementary Figure S2). Hence, these summary statistics distinguish different regulatory mechanisms. Existing models of metabolic labelling protocols are limited to averages across cells as they employ deterministic approaches [26, 27, 49–53, 64] or they are restricted to labelled transcripts [55, 56] and hence cannot make use of dynamic correlations.

Our approach, which is based on the ABC rejection method, effectively samples from the full posterior distribution across parameters and models by comparing millions of candidate simulations to any number of genes in parallel. This is equivalent to recycling all particles for each gene. The method results in an amortised inference using data-independent simulations that is more efficient than data-specific approaches such as sequential Monte Carlo [60]. Amortised simulation-based inference incurs an up-front computational cost, but is applicable to new data points efficiently [65]. Our method is thus ideally suited for the analysis of genome-wide data [66].

A distinctive feature of our approach is that it simultaneously corrects for technical variation through cell-specific capture efficiency and gene-specific metabolic labelling efficiency. Previous inference methods utilising metabolic labelling data relied on specific distribution assumptions [26, 56] or on RNA spike-ins [56] for modelling capture efficiency, while other studies modelled labelling efficiency using either a global rescaling or mixture models [27, 49–51, 54, 64, 67]. We verified that capture efficiency was cell cycle-independent, thus validating that we can distinguish the effects of cell cycle and size. We also showed that labelling efficiency varied dramatically and correlated well with gene length but did not bias our model selection, thus demonstrating that one needs to calibrate labelling efficiency for parameter estimation. The observed correlation between labelling efficiency and gene length may provide insights into the mechanistic basis of metabolic labelling.

By removing technical noise effects, we were able to dissect sources of biological noise contributing to the total transcript variability. By quantifying absolute kinetic rates, we revealed how these sources of transcriptional variation depend on decay rate. We found that mRNA synthesis and degradation kinetics and cell cycle-dependent variation are the primary genome-wide sources of noise, while only a small fraction of the total noise stems from transcriptional bursting. The low levels of bursting noise were explained by overall low burst sizes and relatively high burst frequencies. These observations indicate that transcript abundance is characterised by minimal intrinsic stochasticity, as demonstrated in [11].

We found that most genes scale transcription with cell size. Genes that did not scale were either involved in growth-independent processes, such as cell surface receptor signalling, or in other processes such as proliferation and development. Cell surface and DNA binding proteins do not scale with cell volume and have been previously identified as potential non-scaling genes [68–70]. Size-scaling genes typically rise more rapidly during the cell cycle, have higher degradation rates, and have higher cell cycle variation than non-scaling genes. This suggests that increased synthesis rate with cell size requires higher degradation for cells to achieve homeostasis [30, 71].

Our model predicts waves of regulation across the cell cycle-dependent transcriptome, with kinetic rates peaking at specific cell cycle phases. We observed no step-wise transcriptional changes, suggesting that gene dosage effects mattered less than specific cell cycle regulation of kinetic rates. Genes with kinetics modulating decay rate peaked first in G1, then burst size peaked in S and S/G2, then burst frequency peaked in S phase and the later stages of the cell cycle. A possible explanation of our findings is that global transcription and degradation regulators act in different cell cycle phases and that burst size and frequency modulation is mediated by different transcription factors. Our predictions agree qualitatively with a similar sharp decrease in burst frequency following S phase observed using smFISH [58], and the increased synthesis rates and decreased decay rates following G1 reported in the original scEU-seq study [26]. A mechanistic understanding of this phenomenon would require a gene regulatory network-based approach that is beyond the scope of our study.

Although we expect our inference method to generalise to more complex models and parametric assumptions, it also has several limitations. One limitation is that ABC rejection sampling can lead to weaker accuracy in parameter estimation and broader posterior distributions compared to optimisation-based schemes (see e.g. Supplementary Figure S3). A hybrid method that uses ABC rejection sampling as a first step, followed by an SMC step where the particles for each gene are optimised, would offer a better tradeoff between parallelisation and accuracy [60, 66]. An alternative approach would be to use a synthetic likelihood or integrate ABC with a machine learning method that allows for surrogate modelling [72]. Another limitation is that we assumed deterministic cell cycles across cells, thus a potential improvement would be to account for cell cycle length variability in our moment estimates.

Other studies use modelling to better understand transcription dynamics through mRNA splicing [64, 73–76] including cell cycle-dependent effects [77] and during development [54, 74]. Therefore, splicing data could be used instead of cell cycle reporters in our inference method. Similarly, splicing kinetics could extend our analysis to understand transcription variation along developmental trajectories [54, 73, 74] or identify hidden sources of cellular variation [40].

Our study illustrates how mechanistic modelling and amortised inference can efficiently integrate data across labelled, unlabelled transcripts and cell cycle reporters and also across different pulse and chase protocols. We have demonstrated that mechanistic models can be used to overcome the challenge of technical noise across modalities commonly suffered by multi-view statistical methods [78, 79]. Data integration across conditions, time and modalities is an important challenge for single-cell multi-omics, including epigenome and proteome [80–82], and mechanistic models like ours could provide a versatile tool for integrating these data [83].

In summary, our study reveals global mechanisms of transcription regulation through stochastic modelling of time-resolved single-cell data. Our inference methodology is highly scalable and translates seemingly to complex models of gene regulation. It is thus well suited to uncover complex mechanisms of gene expression control from single-cell transcriptomics. We thus expect that our framework could contribute to uncover regulatory mechanisms that drive cell fate decisions or their dysregulation in disease.

## Data and code availability

Data files that include the accepted particles per model and per gene and all outputs of ABC simulations can be found in the Zenodo repository https://doi.org/10.5281/zenodo.10724157. Julia code used for modelling and data analysis as well as other useful datasets are available in the Github repository https://github.com/pthomaslab/abc_inference_transcription.

## Acknowledgements

This work was supported by UKRI through a Future Leaders Fellowship [MR/T018429/1 to P.T.]; and the Department of Mathematics, Imperial College London through a Roth scholarship [to D.V.]. We would like to thank Nicolas Battich for sharing and discussing their published experimental data from the scEU-seq study. Also, we would like to acknowledge Wenhao Tang, Fern Hughes, Francesco Puccioni, Hannah Dewhurst, Paul Piho and Johannes Pausch for useful conversations.

## Supplementary Methods

In this section, we outline the details of our modelling framework, including data pre-processing and selection of summary statistics, construction of the model, parameter inference and model selection.

### 1.1 Data description, data pre-processing and summary statistics

#### 1.1.1 Unlabelled and labelled mRNA count matrices

The scEU-seq dataset [26] includes count matrices for 4 transcriptomic modalities: unlabelled unspliced (**UU**), unlabelled spliced (**US**), labelled unspliced (**LU**) and labelled spliced (**LS**) mRNA. Our study is focused on unlabelled and labelled mRNA, so we initially add the unspliced and spliced counts to create the (total) unlabelled (**U**) and (total) labelled (**L**) mRNA count matrices: **U** = **UU** + **US, L** = **LU** + **LS**.

#### 1.1.2 Grouping cells with respect to pulse-chase labelling condition

The dataset contains distinct subpopulations of cells that are treated with different metabolic labelling conditions of various durations (*t*_*pulse*_) and various sequencing times after labelling (*t*_*chase*_). These include the *pulse* conditions (*c*_*pulse*_) that involve pulse labelling for time *t*_*pulse*_ as well as the *chase* conditions (*c*_*chase*_) that involve pulse labelling for *t*_*pulse*_ = 22 hours followed by a chase experiment for time *t*_*chase*_. The various combinations of (*t*_*pulse*_, *t*_*chase*_) for each case are outlined in the following tables (time in hours):

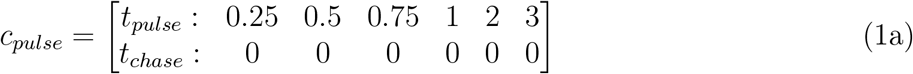

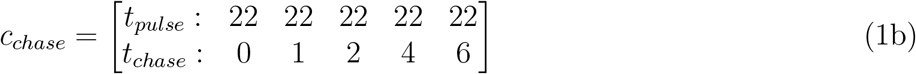

Part of our summary statistics are computed based on pulse-treated and chase-treated cells separately, as there are significant differences between the two groups in the data (see for example Figure 3 (B)). Also, to simplify the notation, we use *t* to denote the dependence of statistics on a specific labelling condition (*t*_*pulse*_, *t*_*chase*_), which we refer to as “labelling time”.

#### 1.1.3 Grouping cells with respect to the cell cycle

The original scEU-seq study provides an estimated cell cycle ordering that assigns each cell to a relative cell cycle position along the interval (0, 1] based on the cell cycle reporters. We rely on these estimates and divide the 5422 cells into 5 groups by partitioning this ordering trajectory into 5 sub-intervals (0, 1*/*5], (1*/*5, 2*/*5], …, (4*/*5, 1]. According to the fluorescence of cell cycle reporters (Supplementary Figure S1), we assume that the first two intervals correspond to G1-G1/S phase, the next two intervals correspond to S-S/G2 phase and the last interval corresponds to G2/M phase. We then use the proportion of cells across cell cycle phases to define an empirical *age distribution* (see Figure 3 (B)):

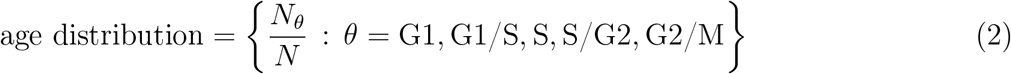

where *N*_*θ*_ denotes the number of cells in cell cycle phase *θ* and *N* = 5422. Additionally, each cell cycle phase includes cells from different labelling conditions (Supplementary Methods 1.1.2), and therefore we also define age distributions per labelling time, 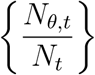, where *N_t_* is the number of cells in condition *t* and *N*_*θ*,*t*_ is the number of cells in condition *t* and cell cycle phase *θ*.

#### 1.1.4 Gene filtering

We impose certain criteria to select genes for parameter inference and downstream analysis. Starting from 11848 genes of the scEU-seq dataset, we first select genes with at least 500 labelled mRNA counts across all cells (similarly to the original study [26]), which leaves us with a subset of 6087 genes. Then, we impose an additional criterion to filter out genes with very low expression. We require that each gene has at least 1 count on average in at least one cell cycle phase. That is, if **X** = (*x*_*ij*_) = **U** + **L** is the total (unlabelled + labelled) mRNA count matrix (where cells are indexed with *i* and genes are indexed with *j*), then the gene-specific condition is that

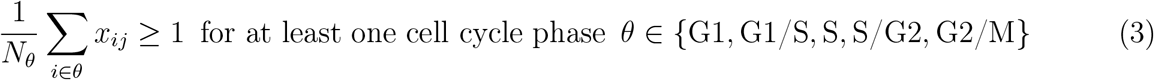

This criterion gives us the final set of 3419 genes used for further analysis.

#### 1.1.5 Summary statistics

We now describe how we calculate summary statistics of data. We consider the count matrices **U** = (*u*_*ij*_), **L** = (*l*_*ij*_) of unlabelled and labelled mRNA, as well as the count matrix **X** = (*x*_*ij*_) = **U**+**L** of total mRNA. The summary statistics of a gene *j* in the data depend on the cell cycle phase *θ* ∈ *{*G1, G1/S, S, S/G2, G2/M*}* or on the labelling condition *c* ∈ *{c*_*pulse*_, *c*_*chase*_*}*. The definitions of gene-specific summary statistics are outlined in Table 1. In these definitions, the notation *i* ∈ *c* refers to either all *pulse*-treated cells or all *chase*-treated cells, while the notation *i* ∈ *t* refers to a specific labelling condition (*t*_*pulse*_, *t*_*chase*_), where *t* = *t*_*pulse*_ + *t*_*chase*_ denotes the total duration of an experiment (see paragraph 1.1.2).

#### 1.1.6 Standard errors of summary statistics

Aside from computing summary statistics from data, we also estimate standard errors for each statistic to quantify the measurement bias. If *s*_obs_ = *s*_obs_(*y*) is a summary statistic of the data *y*, we estimate the standard error of *s*_obs_ with

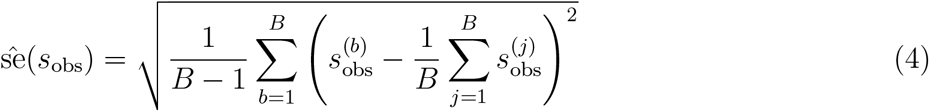

where 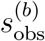 is the value of the summary statistic that we obtain from each bootstrapped dataset *y*^(*b*)^, *b* = 1, …, *B* = 100. These standard errors are used during parameter inference to normalise the magnitudes of different summary statistics (see Supplementary Methods 1.5.1).

### 1.2 Stochastic modelling framework

We develop a stochastic model that effectively describes the underlying generative process of mRNA transcription in cells growing and dividing and going through the scEU-seq labelling protocols.

#### 1.2.1 Model of bursty transcription of unlabelled and labelled mRNA

We assume that mRNA molecules are produced according to a telegraph model of gene expression, capturing random switching of the gene promoter between active and inactive states with rates *k*_on_ and *k*_off_ respectively, mRNA synthesis with rate *α* and mRNA decay with rate *γ*. During pulse-chase labelling, labelled and unlabelled mRNA molecules are synthesised with rates *λ α* and (1 − *λ*) *α* respectively, where *λ* denotes the gene-specific labelling efficiency. These are encoded in the reaction scheme:

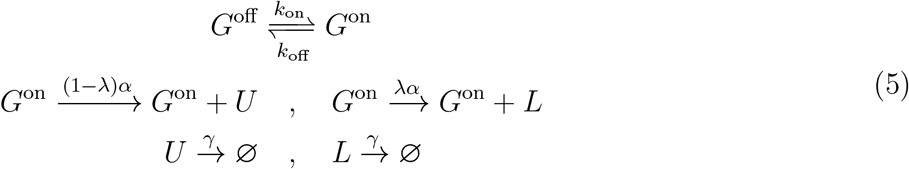

#### 1.2.2 Cell cycle-dependent kinetic rates

We further allow for three of these kinetic rates to vary over the cell cycle. As a result, they are time-dependent periodic functions *k*_on_ = *k*_on_(*t*), *α* = *α*(*t*), *γ* = *γ*(*t*), for 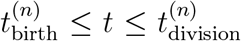, *n* = 1, 2, … (with *n* representing the *n*^th^ cell generation in a single-cell lineage). As they are periodic functions, their value does not depend on *n* and their cell cycle-dependence can be equivalently written as

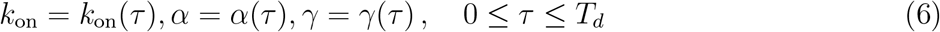

where *τ* is the cell age (time along cell cycle progression) and 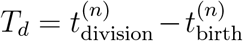 is the interdivision time (cell cycle length). The rate *k*_off_ is always assumed to be constant. An additional assumption on the kinetics is that the synthesis rate *α* scales linearly with cell size (except for the constant non-scaling model). This is in accordance with observations from the data S1, which allows us to assume that cells follow linear growth. This implies that the cell size *s* at age *τ* is:

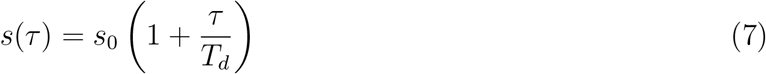

where *s*_0_ is the cell size at birth. Accordingly, we model the dependence of synthesis rate on cell size as:

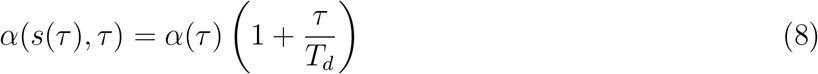

where *α*(*τ*) describes the dependence of the synthesis rate on cell cycle progression without size-scaling. Additionally, we define the burst kinetics as follows:

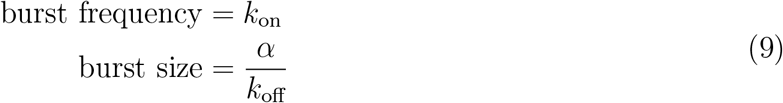

As a consequence, both burst frequency and burst size can be cell cycle-dependent and additionally the burst size could be size-dependent (except for the constant non-scaling model), in accordance to the synthesis rate. The unit of all kinetic parameters is *h*^−1^, apart from burst size which is unitless. Finally, we define the mRNA half-life *t*_1*/*2_ (measured in hours) that corresponds to a constant degradation rate *γ* as:

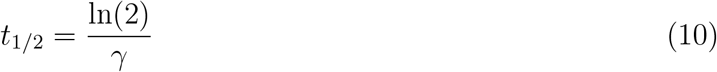

#### 1.2.3 Moment equations with time-dependent kinetic rates

The joint probability distribution of molecule numbers **x** = (*x*_1_, *x*_2_, *x*_3_)^T^ of the respective species *{G*^on^, *U, L}* in the reaction scheme (5) obeys the Chemical Master Equation (CME):

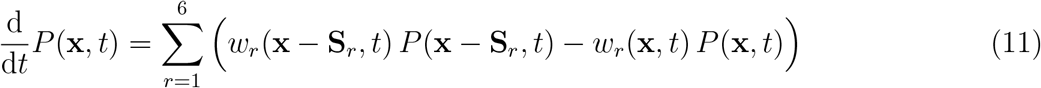

where **S**_*r*_ = (**S**_1*r*_, **S**_2*r*_, **S**_3*r*_)^T^ denotes the *r*-th column of the stoichiometric matrix

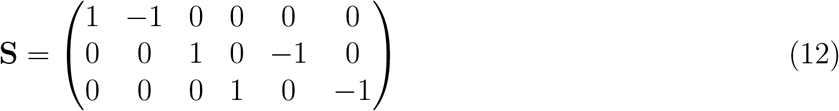

of the reaction system (with **S**_*ir*_ being the change in molecule number of species *i* by reaction *r*) and *w*_*r*_(**x**, *t*) denotes the propensity of reaction *r*, which is the probability per unit time for the *r*-th reaction to occur. The propensities of the network are

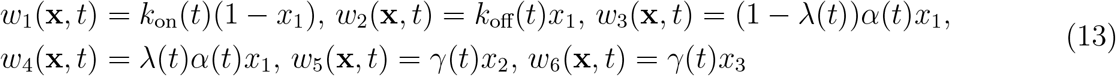

Note that the gene promoter state *x*_1_ is either 0 or 1 so it is sufficient to only describe the state of *G*^on^ in the equations. In our model, we are interested in the time-evolution of the 1^st^ and 2^nd^ order central moments of the molecule abundances, which can be derived using a moment-closure approximation of the CME [38]:

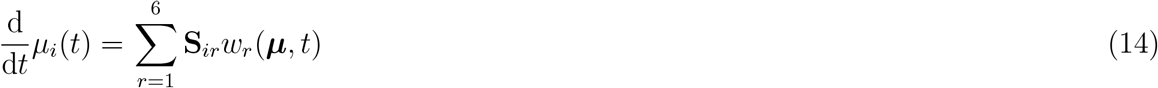

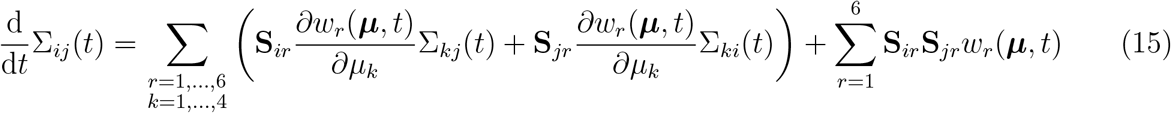

where *µ*_*i*_ = 𝔼 (*x*_*i*_) is the mean molecule number of species *i* and **Σ** = (Σ_*ij*_) is the covariance matrix with Σ_*ij*_ = 𝔼 ((*x*_*i*_ − *µ*_*i*_)(*x*_*j*_ − *µ*_*j*_)) being the covariance of molecule numbers of species *i* and *j* and the diagonal elements Σ_*ii*_ = Var(*x*_*i*_) being the variance of molecule number of species *i*. Note that since the reactions in (5) are linear, the propensities are linear functions of the abundances *x*_*i*_, that is 𝔼 (*w*_*r*_(**x**, *t*)) = *w*_*r*_(𝔼 (**x**), *t*) = *w*_*r*_(***µ***, *t*), and therefore the system of moment equations is closed (does not depend on higher order moments). Given our assumption about cell cycle-dependence of the reaction rate constants, the propensities are periodic functions of time. As a consequence, the moment equations are constrained by periodic boundary conditions.

#### 1.2.4 Coupling to cell division dynamics

To determine the boundary conditions, we model the random partitioning of molecules at cell division as a binomial process, which gives us the following conditions for the mean, variance and covariance of unlabelled and labelled mRNA at the *n*-th cell division:

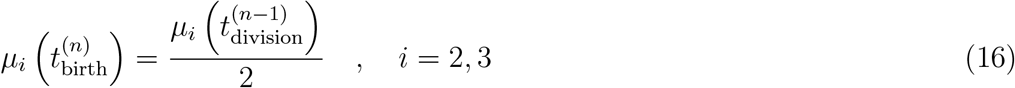

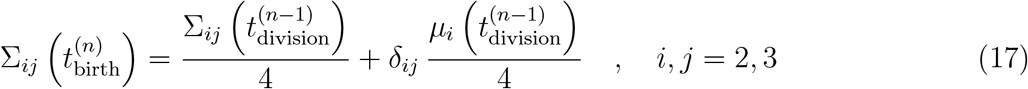

The gene promoter state *g*^(birth)^ in a daughter cell at birth is identical to the promoter state *g*^(division)^ in the mother cell at division, and therefore the mean and variance of the promoter state remain constant before and after division:

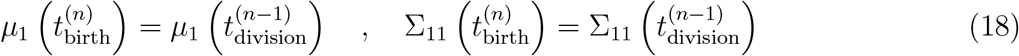

However, the covariance of the promoter state with the mRNA decreases due to partitioning of mRNA molecules:

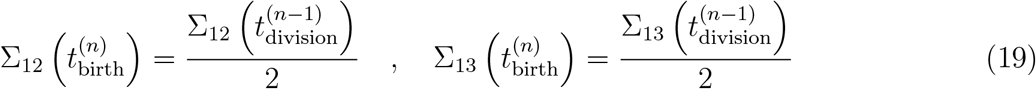

To prove the above, for notation simplicity, *m* denotes mRNA (unlabelled or labelled) and *g* denotes the gene promoter state. Also, *m*_0_, *g*_0_ denote the mRNA and gene states in a daughter cell at birth and *m*_*T*_, *g*_*T*_ denote the mRNA and gene states in the mother cell at cell division. At birth, a daughter cell inherits *m*_0_ mRNA molecules out of total *m*_*T*_ molecules of the mother cell according to a Binomial distribution:

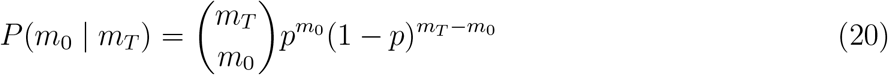

with *p* = 1*/*2. Also, the gene promoter state is not altered at cell division and therefore:

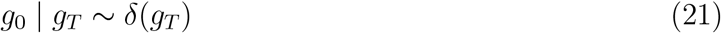

Using the law of total expectation, the expected number of mRNA at cell birth is:

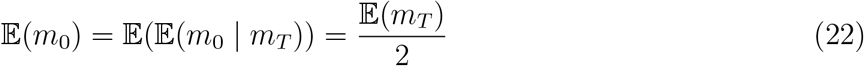

Using the law of total variance, the variance of mRNA molecules at birth is:

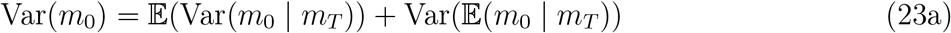

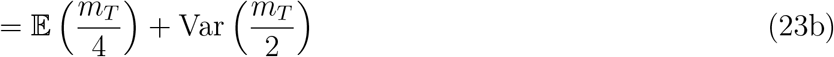

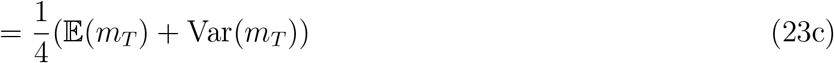

Denoting with *u, l* the unlabelled and labelled mRNA and using the law of total covariance, we have that the covariance of unlabelled and labelled mRNA at birth is:

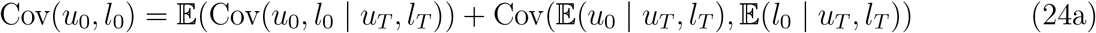

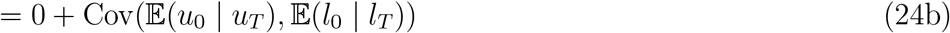

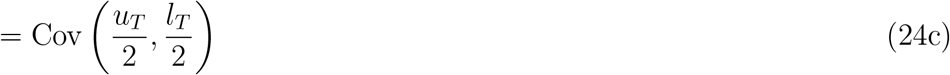

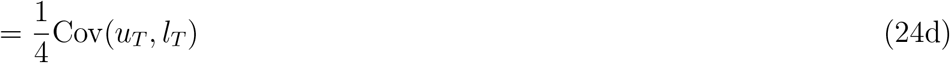

where in (24b) we use that the conditional variables *u*_0_ | *u*_*T*_ and *l*_0_ | *l*_*T*_ are independent.

The covariance of the mRNA and gene states at birth is given by:

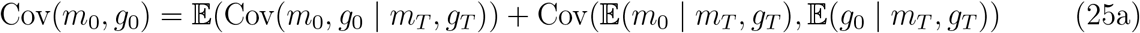

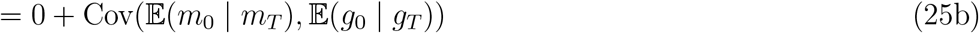

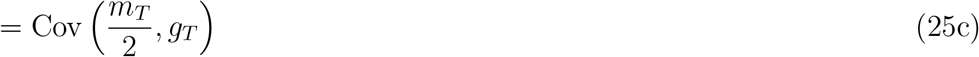

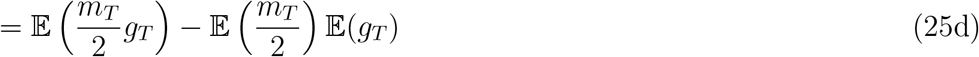

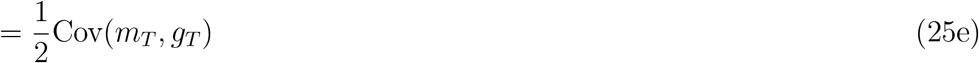

where in (25b) we use that the conditional variables *m*_0_ | *m*_*T*_ and *g*_0_ | *g*_*T*_ are independent.

#### 1.2.5 Partitioning of 3rd order moments

In the case where we want to extend the moment equations of the model up to 3rd order, we need additional formulas for partitioning the 3rd order central moments. Starting from the 3rd central moment of the mRNA, we use the law of total cumulance to get:

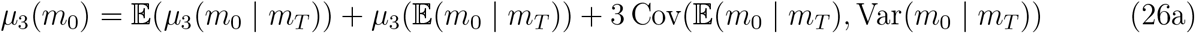

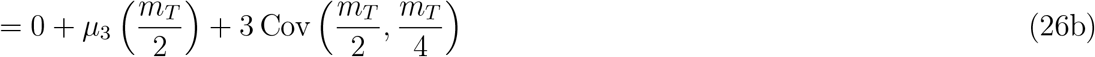

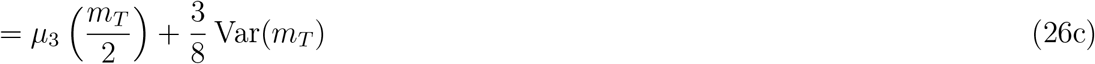

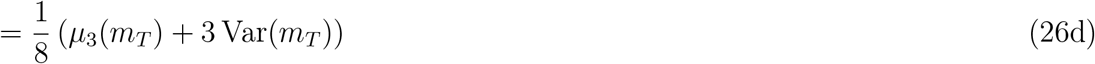

Similarly, for the 3rd central moment of the gene promoter:

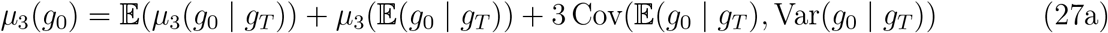

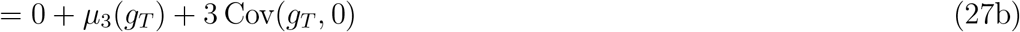

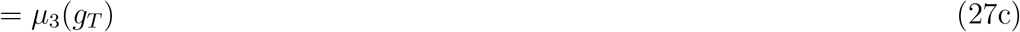

Finally, we derive the partitioning for the mixed 3rd central moments of the gene promoter and the mRNA. We get that:

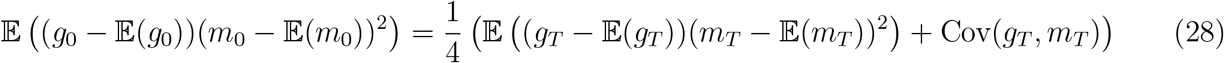

and

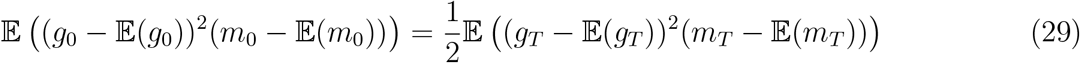

where we are using the following formulas:

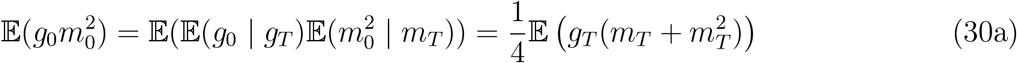

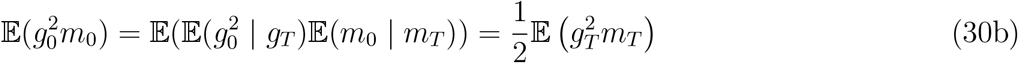

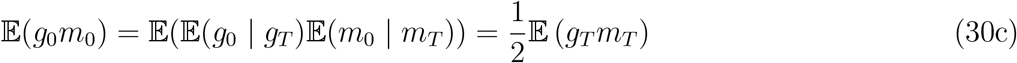

#### 1.2.6 Description of a single realisation of the model

A single model realisation depends on a gene-specific parameter set *{k*_off_, *k*_on_(*t*), *α*(*t*), *γ*(*t*), *λ}*, a cell age *τ* and a labelling condition (*t*_*pulse*_, *t*_*chase*_). We solve the moment equations numerically over time for multiple consecutive cell divisions, following a single-cell lineage up until the sequencing time-point *t*_*seq*_ (see example model realisations in Figure 1 (C) and Supplementary Figure S2). We additionally assume that cells have a deterministic cell cycle length, *T*_*d*_ = 20 hours. First, the equations are solved for multiple cell cycles until convergence to a steady state. The convergence criterion is that the relative change of all trajectories at the end of two consecutive cell cycles needs to be smaller than 0.01%. After reaching convergence, we solve the model for additional time equal to 3 *T*_*d*_ + *τ*, up until the final time-point *t*_*seq*_ (observation or sequencing time-point). To model the effect of pulse labelling on mRNA synthesis, the labelling efficiency *λ*(*t*) is set to its constant parameter value *λ* for time *t*_*pulse*_:

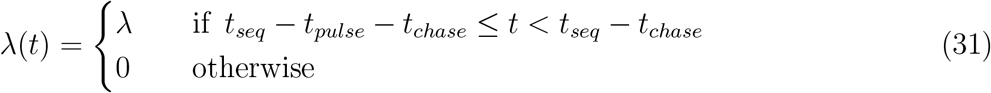

The endpoints 𝔼 (*u*), 𝔼 (*l*), Var(*u*), Var(*l*), Cov(*u, l*) of the trajectories at time *t*_*seq*_ comprise the model estimates, which are conditional on cell age *τ* and pulse-chase labelling time (*t*_*pulse*_, *t*_*chase*_).

### 1.3 Capture efficiency estimation and downsampling of moments

To take into account technical effects of sequencing in our modelling framework, we correct the model outputs using estimates of cell-specific capture efficiencies. Consider the count matrices **U** = (*u*_*ij*_) and **L** = (*l*_*ij*_) of unlabelled and labelled mRNA, as well as the total mRNA count matrix **X** = (*x*_*ij*_) = **U** + **L**, where the indices *i* = 1, …, *n* represent cells and the indices *j* = 1, …, *d* represent genes. If *u* = *u*_*ij*_ and *l* = *l*_*ij*_ are a pair of observations for a cell *i* and gene *j*, then due to low transcript capture of the sequencing protocol, these observations are random samples drawn from Binomial distributions, *u* ∼ Binomial 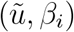 and *l* ∼ Binomial 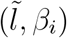, where 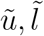 are the mRNA counts we would observe in a sequencing experiment with 100% capture efficiency and *β*_*i*_ is the capture efficiency of cell *i*. Below, we outline the method for estimating the capture efficiency distribution directly from the data and provide formulas for the corrected (downsampled) moments of *u* and *l*.

#### 1.3.1 Estimation of cell-specific capture efficiencies from scEU-seq data

The estimation of *β*_*i*_ is based on the assumption that cells in the same cell cycle phase have on average the same cell size. We use the total mRNA abundance 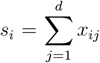 of cell *i* of as a proxy for its size, while the average total mRNA count 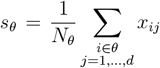 of cells in cell cycle phase *θ* represents the characteristic cell size in *θ* (*N*_*θ*_ is the number of cells in *θ*). Based on the above, we can estimate:

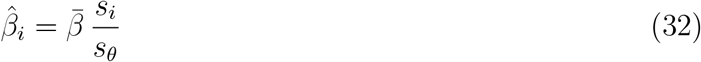

where 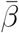 is an estimate of the average capture efficiency of the sequencing experiment.

An additional factor that we take into account in the above estimation is that cells treated in pulse conditions have a different capture efficiency than cells treated in chase conditions (see Figure 3 (B) and Supplementary Methods 1.1.2). For that reason, we consider different average capture efficiencies 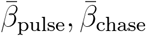 for pulse-treated and chase-treated cells, and the normalising factors *s*_*θ*_ for each cell cycle phase are evaluated independently for pulse and chase cells. The estimated capture efficiency 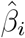 of cell *i* is hen given by:

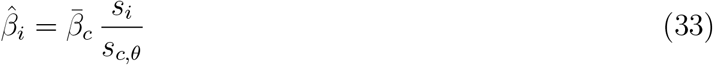

where 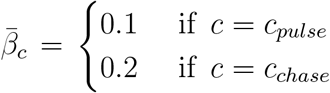 is the average capture efficiency of pulse/chase experiments and 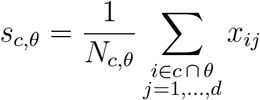 is the average total count of cells in condition *c* ∈ *{c*_*pulse*_, *c*_*chase*_*}* cell cycle phase *θ*. The ratio of the values we have assumed for 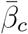 (capture efficiency of chase being twice the capture efficiency of pulse) is inferred from the data, while the absolute scale of 10% capture efficiency is an estimate that is reasonable for scRNA-seq protocols [47], but it is not typically directly measured.

#### 1.3.2 Downsampling the mean, variance and covariance

We can now use the estimated capture efficiencies to incorporate the effect of Binomial downsampling of data in the moments. For a cell with capture efficiency *β*, the downsampled mean mRNA level is given by:

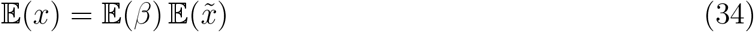

where 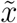 is the mRNA levels in the absence of technical noise. The downsampled variance is given by:

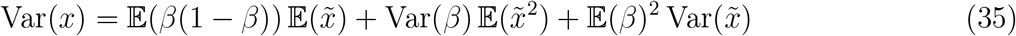

The formulas (34),(35) hold for both unlabelled and labelled mRNA.

Finally, the downsampled covariance of unlabelled and labelled mRNA is given by:

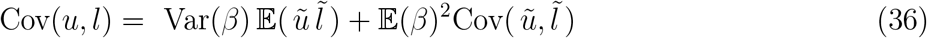

To prove the above, we use that an observed mRNA count *x*_*ij*_ = *x* is a random variable *x* ∼ Binomial(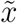, *β*) where *β* = *β*_*i*_ is the capture efficiency of cell *i* and 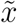 is the observation in the absence of technical errors. The downsampled mean is given by:

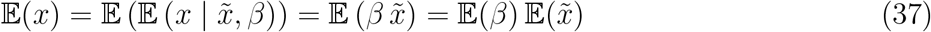

and the downsampled variance is given by:

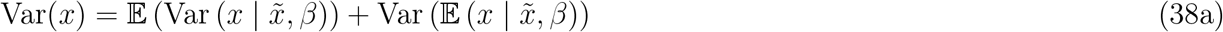

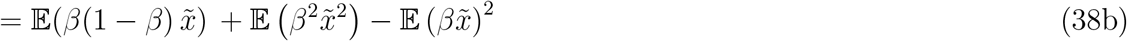

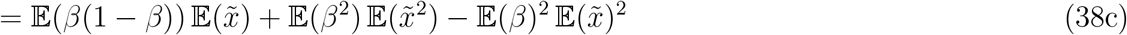

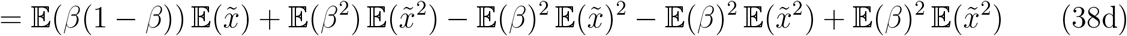

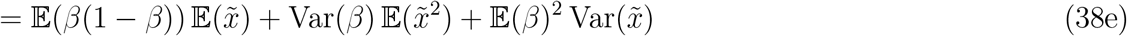

Finally, the downsampled covariance of unlabelled and labelled mRNA is given by:

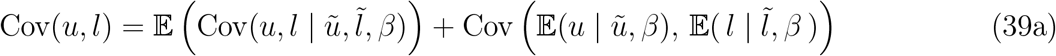

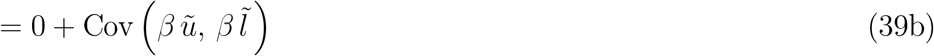

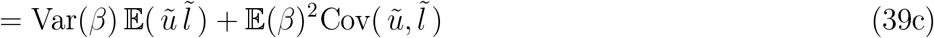

### 1.3.3 Downsampling the 3rd central moment

Following similar principles as above, we derive the binomial downsampling of the 3rd order central moment. We here only consider the 3rd moment of the total mRNA count *x* = *u* + *l* and not the mixed 3rd order moments. Using the law of total cumulance, we get:

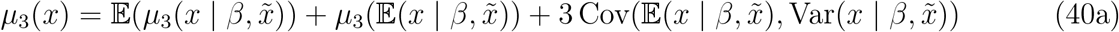

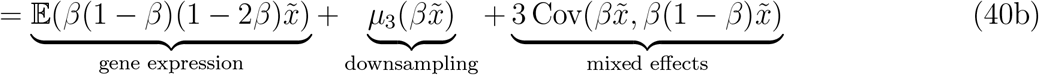

where

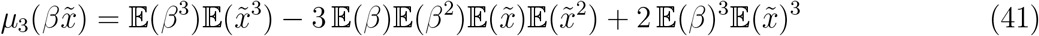

and

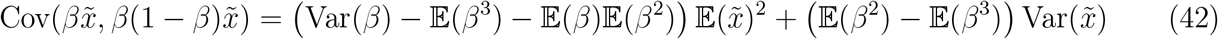

### 1.4 Time-dependent summary statistics generated from the model

We have demonstrated how our model generates 1^st^ and 2^nd^ order moments that are dependent on cell age *τ* and labelling time *t* (Supplementary Methods 1.2.6), as well as how we correct them to account for technical noise (Supplementary Methods 1.3.2). Below, we describe how we further use these moments to construct a collection of summary statistics and extract interpretable information about bursty transcription dynamics.

### 1.4.1 Cell cycle-dependent mean and noise statistics

First, we construct summary statistics that describe cell cycle-dependence of mean and noise in transcript levels. Given the means of unlabelled and labelled mRNA 𝔼 (*u* | *τ, t*), 𝔼(*l* | *τ, t*), we compute the mean of total transcript levels *x* = *u* + *l*:

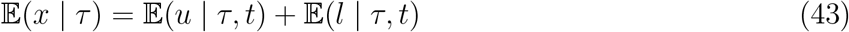

Note that this mean does not depend on labelling condition *t*. Similarly, we compute the variance of total transcripts 2^nd^ order moments of unlabelled and labelled mRNA:

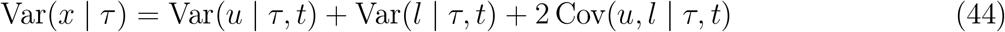

as well as the Fano factor:

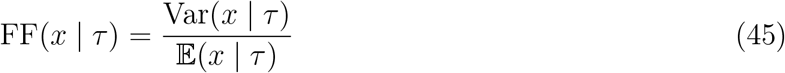

These summary statistics describe temporal changes of transcription dynamics along the cell cycle that are independent of variations due to labelling.

#### 1.4.2 Labelled vs total mRNA ratios capture the timescales of RNA kinetics

Next, we generate summary statistics that aim to extract information about the timescales of bursty transcription and degradation from the various pulse and chase labelling conditions. We first construct a statistic that describes how the proportion of labelled mRNA changes with labelling time:

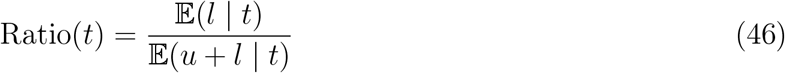

To integrate out the cell cycle-dependence we use empirical age distributions from the data, which depend on the labelling condition (Supplementary Methods 1.1.3):

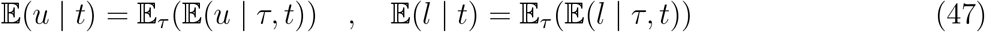

#### 1.4.3 Labelled-unlabelled mRNA correlations capture sources of transcriptional noise

Further, we construct summary statistics that describe the covariation of labelled and unlabelled mRNA. The law of total covariance gives:

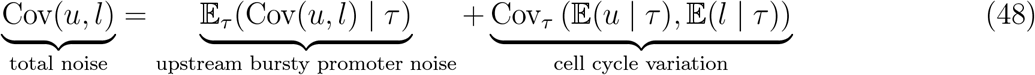

which shows that the unlabelled-labelled RNA modalities and the cell cycle reporter provided by the scEU-seq protocol can be used to decompose the overall noise into two contributions: upstream noise induced by the bursty gene promoter and noise due to dependence on the cell cycle (see also Figure 3 (A)). To exploit this information, our summary statistics include two correlation coefficients - corresponding to the two noise components - that contribute to the total correlation of unlabelled-labelled mRNA:

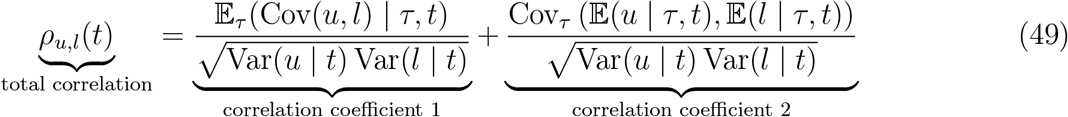

Similarly to the ratios (Supplementary Methods 1.4.2), we compute labelling-specific variances and covariances by integrating over the labelling-specific age distributions. Note that the total variances in the denominators of (49) are given by the law of total variance:

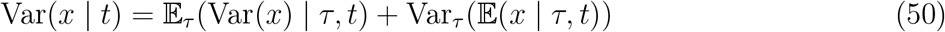

### 1.5 Approximate Bayesian inference framework

In this section, we outline the details of our inference framework. As our moment-based model does not provide us with an analytical likelihood, relying on numerical solutions of our model, we employ an Approximate Bayesian approach for estimating parameters, where generated summary statistics from the model are compared with summary statistics of the data.

#### 1.5.1 ABC rejection sampling for parameter inference

For each gene *j* in the data, we consider its associated collection of summary statistics 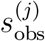 (as de-scribed in Supplementary Methods 1.1.5, 1.1.6). The scheme for estimating gene-specific parameters is the following:

For each model index *m* = 1, …, 5 and for *i* = 1, …, *M*,

- Sample a parameter set *θ*^(*i*)^ = *θ*^(*i*)^(*m*) from the prior *π*(*θ*).
- Generate synthetic summary statistics *s*^(*i*)^ = *s*^(*i*)^(*θ*^(*i*)^) using the model.
- Compute distances *d*(*s*^(*i*)^, 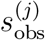) for all *j* = 1, …, *N* .
- Accept parameter set *θ*^(*i*)^ for all *j* = 1, …, *N* that satisfy *d*(*s*^(*i*)^, 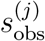) ≤ *ε*, where *ε* is a predetermined acceptance threshold.

For each *j* = 1, …, *N*, the collection of accepted parameter sets *{*Θ^(*j*)^(*m*) : *m* = 1, …, 5*}* comprises an approximate (joint) posterior distribution over parameters and models for gene *j*. After model selection (see Supplementary Methods 1.5.2), we choose the best model *m*^⋆^ = *m*^⋆^(*j*) for each gene *j* and we use the particle *θ*^(*i*)^(*m*^⋆^(*j*)) that achieved the minimum distance *d*(*s*^(*i*)^, 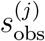)^⋆^ between model and data as the gene-specific point estimate to inform the model for downstream analyses.

To determine the prior distribution *π*(*θ*), we sample parameters from the following log-uniform distributions: log_10_(*k*_on_) ∼ *𝒰* (−3, 3), log_10_(*k*_off_) ∼ *𝒰* (−3, 3), log_10_(*α*) ∼ *𝒰* (−3, 3), log_10_(*γ*) ∼ *𝒰* (−3, 2) and log_10_(*λ*) ∼ *𝒰* (−0.7, 0). We sample *M* = 10^6^ parameter sets for each model *m* (Figure 2(A)) so we run in total 5 *·* 10^6^ simulations (this is the computationally expensive step but is independent of the specific data for individual genes hence making our method scalable). For each sampled parameter set *θ*^(*i*)^ = *θ*^(*i*)^(*m*), we generate summary statistics *s*^(*i*)^(*θ*^(*i*)^) from the model *m* according to the procedure described in Supplementary Methods 1.3, 1.4, which we then compare to all *N* = 3419 gene-specific sets of summary statistics. The distance between a synthetic and an observed collection of summary statistics, *s* = *{s*_1_, …, *s*_*n*_*}* and *s*_obs_ = *{s*_obs,1_, …, *s*_obs,*n*_*}*, is determined by the following function:

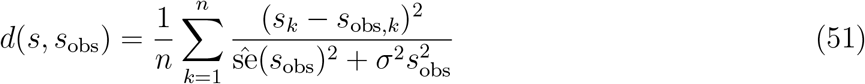

where *n* = 53, 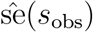 is the standard error of the summary statistic due to sampling noise (Supplementary Methods 1.1.6) and *σ* = 0.1 models other technical variations.

#### 1.5.2 ABC-based model selection

As we are testing multiple modelling hypotheses against all genes in the data (Figure 1 (C), Figure 2 (A)), we wish to assign probabilities to these hypotheses and decide which model is most likely to describe each gene. To do that, we use an ABC-based model selection approach [61], which involves two decision steps. The first one is to select whether a gene is constant or non-constant (see Figure 2 (A), Figure 2 (B)). Given this decision, the second step is to select a specific constant model (scaling, non-scaling) or a specific non-constant model (varying burst frequency, varying burst size or varying decay rate). Let *H*_1_, *H*_2_ denote the constant and non-constant model hypotheses and let *𝒟* denote the data. As we have in total two constant models and three non-constant models, we set the prior probabilities of the two hypotheses to be 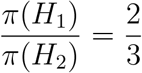. The posterior probability of model *H*_*i*_ is approximated by:

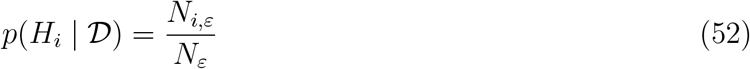

where *N*_*i*,*ε*_ is the number of particles accepted for model *H*_*i*_ and *N*_*ε*_ is the total number of particles accepted for all models, given the acceptance threshold *ε*. In this case, *N*_1,*ε*_ includes the number of particles accepted for both constant models, while *N*_2,*ε*_ includes the number of particles accepted for all three non-constant models. Model comparison is then based on the (approximate) Bayes factor

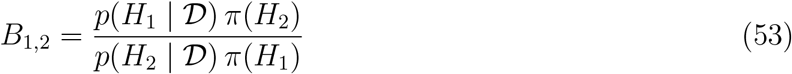

We then need a criterion for thresholding this Bayes factor to decide whether we have sufficient evidence in favour of *H*_1_ over *H*_2_, to a certain level of confidence. To do that we use bootstrapping of the accepted particles for each model, and estimate confidence intervals 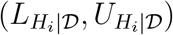 for each posterior probability, so that

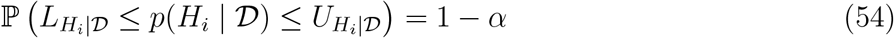

with *α* = 0.05. For model *H*_1_ to be selected versus model *H*_2_ it is required that

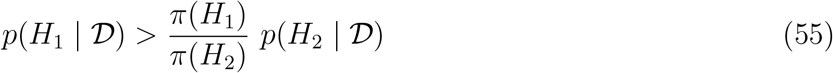

or equivalently, since *p*(*H*_2_ | *𝒟*) = 1 − *p*(*H*_1_ | *𝒟*), we need

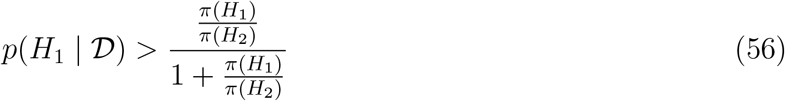

To strengthen this criterion, we require

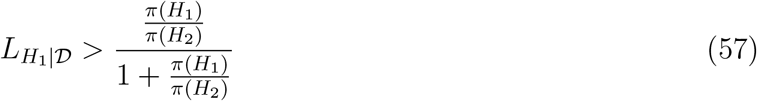

Equivalently, model *M*_2_ is selected if

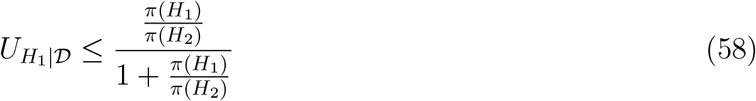

After selecting between the constant and non-constant hypotheses, we proceed to the second step of model selection, where we follow a similar approach. If the constant hypothesis is selected, we now decide between the scaling *H*_1_ and non-scaling *H*_2_ hypotheses, with probabilities given by (52), where now *N*_1,*ε*_, *N*_2,*ε*_ are the number of accepted particles of the scaling and the non-scaling model respectively and *N*_*ε*_ = *N*_1,*ε*_ + *N*_2,*ε*_. If the non-constant hypothesis is selected, we decide between the varying burst frequency *H*_1_, varying burst size *H*_2_ and varying decay rate *H*_3_ hypotheses. In that case, the choice criterion is formed as follows. The model probabilities *p*(*H*_*i*_ | *D*) are given by (52), where now *N*_*i*,*ε*_ is the number of accepted particles of each non-constant model and *N*_*ε*_ = *N*_1,*ε*_ + *N*_2,*ε*_ + *N*_3,*ε*_. If 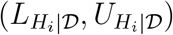 are the corresponding confidence intervals from bootstrapping, we select the non-constant model *i* ∈ *{*1, 2, 3*}* if

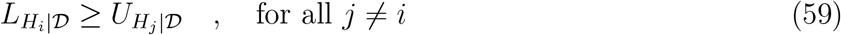

### 1.6 Testing the model performance on synthetic data

We generated synthetic data to test the robustness of our inference and model selection schemes. Using the Gillespie algorithm we performed stochastic simulations of the gene expression model (5) with reaction propensities (13). For each single cell and each gene, we generated unlabelled and labelled transcript counts by sampling from the respective stochastic trajectories of transcript levels at steady state. We used 15*5 ground truth parameters sampled from the posterior distributions we inferred from real data, and simulated 15 genes from each of the 5 model classes (Figure 2 (A)). For each gene, we simulated a total of 2463 cells, across the 11 pulse-chase labelling conditions and the 5 cell cycle stages (Supplementary Methods 1.1.2 and 1.1.3). The distribution of cells across labelling conditions and cell cycle stages corresponded to the one observed in the real scEU-seq data. Additionally, we added technical noise to the simulated count data using Binomial downsampling and cell-specific capture efficiencies that were sampled from Beta distributions. These Beta distributions were chosen by fitting the empirical capture efficiency distributions of pulse and chase cells that we previously estimated from the real data (Supplementary Methods 1.3.1).

We performed inference on the simulated data, followed by model selection, using the framework we described in Supplementary Methods 1.5. For comparison purposes, we refer to this as the *full model*. We next performed inference on 3 additional scenarios:

- Small number of cells: We used half of the cells in the synthetic data (N = 1232) to compute summary statistics.
- Small number of prior samples: We used 2 * 10^5^ prior parameter sets from each model to fit the data (1/5 of the parameters used in the full model).
- Narrow priors: In particular, the parameters used for inference where samples of the following log-uniform distributions: log_10_(*k*_on_) ∼ *𝒰* (−2.5, 2.5), log_10_(*k*_off_) ∼ *𝒰* (−2.5, 2.5), log_10_(*α*) ∼ *𝒰* (−2.5, 2.5), log_10_(*γ*) ∼ *𝒰* (−2.75, 1.75) and log_10_(*λ*) ∼ *𝒰* (−0.6, −0.1).

### 1.7 Noise decomposition

Let *x* be the mRNA count of a gene and *u, l* the corresponding unlabelled and labelled modalities, so that *x* = *u* + *l*. Also, let *τ* denote the cell age and *t* the labelling time (based on experimental condition). Then, using the conditional moments of the model output, we compute the mean transcript levels as:

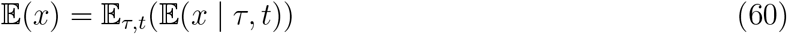

Then, the total variance of *x* can be decomposed as follows:

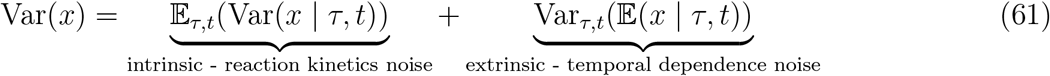

The intrinsic noise component can be further decomposed as follows:

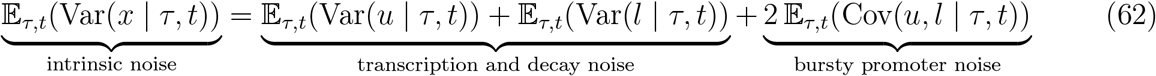

where the transcription and decay term includes noise due to random partitioning at cell division. The extrinsic noise component is explained by variations due to cell cycle-dependence and labelling time:

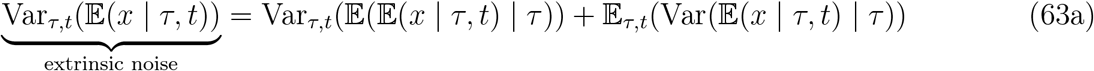

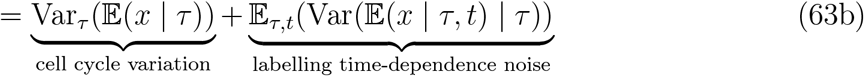

Computing the labelling time-dependence component from the model yields a negligible, practically zero contribution and therefore we can neglect it from the formulas.

We finally obtain the following total variance decomposition:

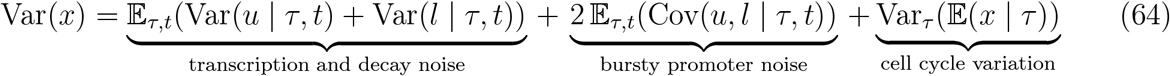

Dividing the whole expression in (64) by the squared total mean E(*x*)^2^, we can obtain the formula for decomposing the total noise 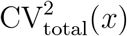. It is important to note that we use the same formula for decomposing the observed total variance in the data and for quantifying cell cycle variation (Figure 4 (E)), however in that case the formula is not exact as it neglects technical noise contributions.

#### 1.7.1 Noise decomposition in the presence of technical noise

We now want to include the effect of technical noise due to cell-specific downsampling in the above decomposition (64). For each individual term in (64), we obtain a correction that includes the effects of technical noise. To correct the transcription and decay noise term, we use the variance downsampling formula (35) to get (for both *x* ∈ *{u, l}*):

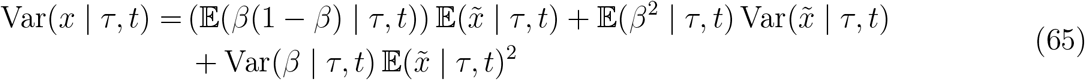

and hence

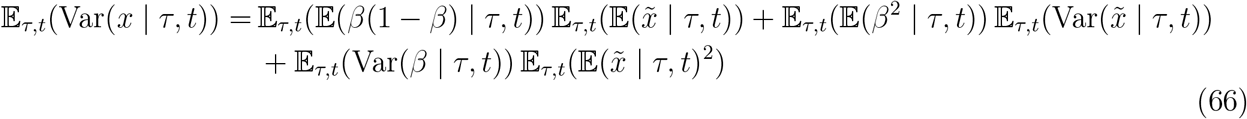

Similarly, to correct the bursty promoter noise term, we use the covariance downsampling formula (36) to get:

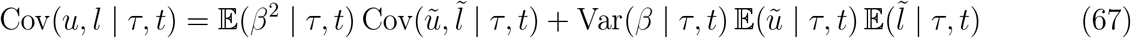

and hence

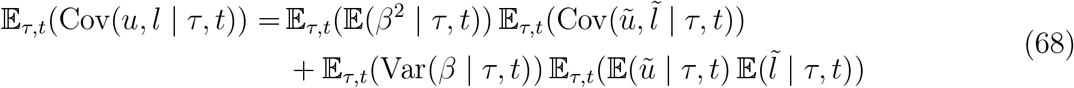

Finally, in the case of the cell cycle variation we use the downsampling of the mean (34):

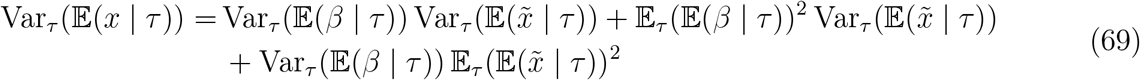

Putting it all together, we get the total variance decomposition formula in the presence of technical noise:

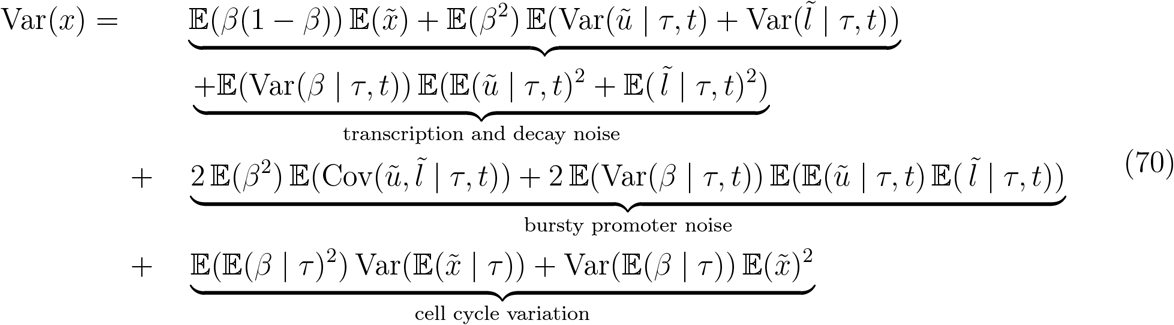

where we have dropped the indices *τ, t* from the total expectations and covariances for notation simplicity.

## Supplementary Figures

**Supplementary Figure S1:**
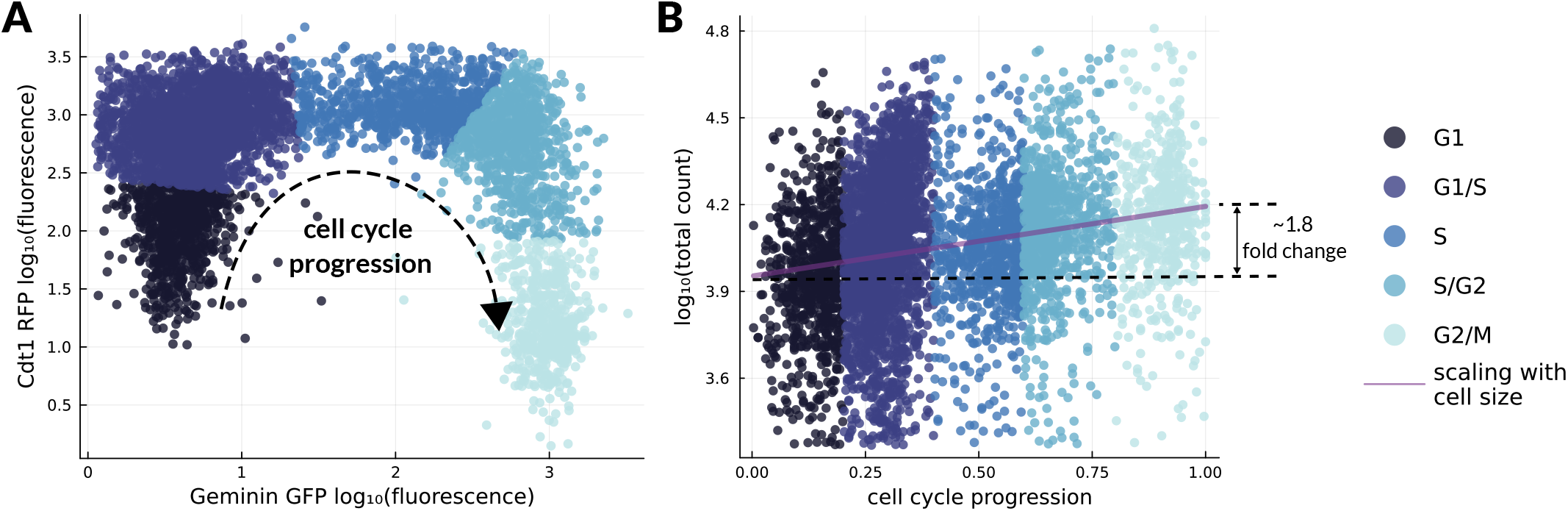
Definition of cell cycle phases and observed scaling of total count with cell size. (A) Scatter plot of log-fluorescence of the cell cycle markers Geminin-GFP and Cdt1-RFP as reported in the original scEU-seq study [26]. The colours show our grouping of cells in cell cycle phases (see also Supplementary Methods 1.1.3). (B) Observed cell-specific total mRNA counts along cell cycle progression, with respect to the cell cycle trajectory reported in [26]. The purple line shows a linear regression fit and the almost 2-fold increase indicates the scaling of total count with cell size.

**Supplementary Figure S2:**
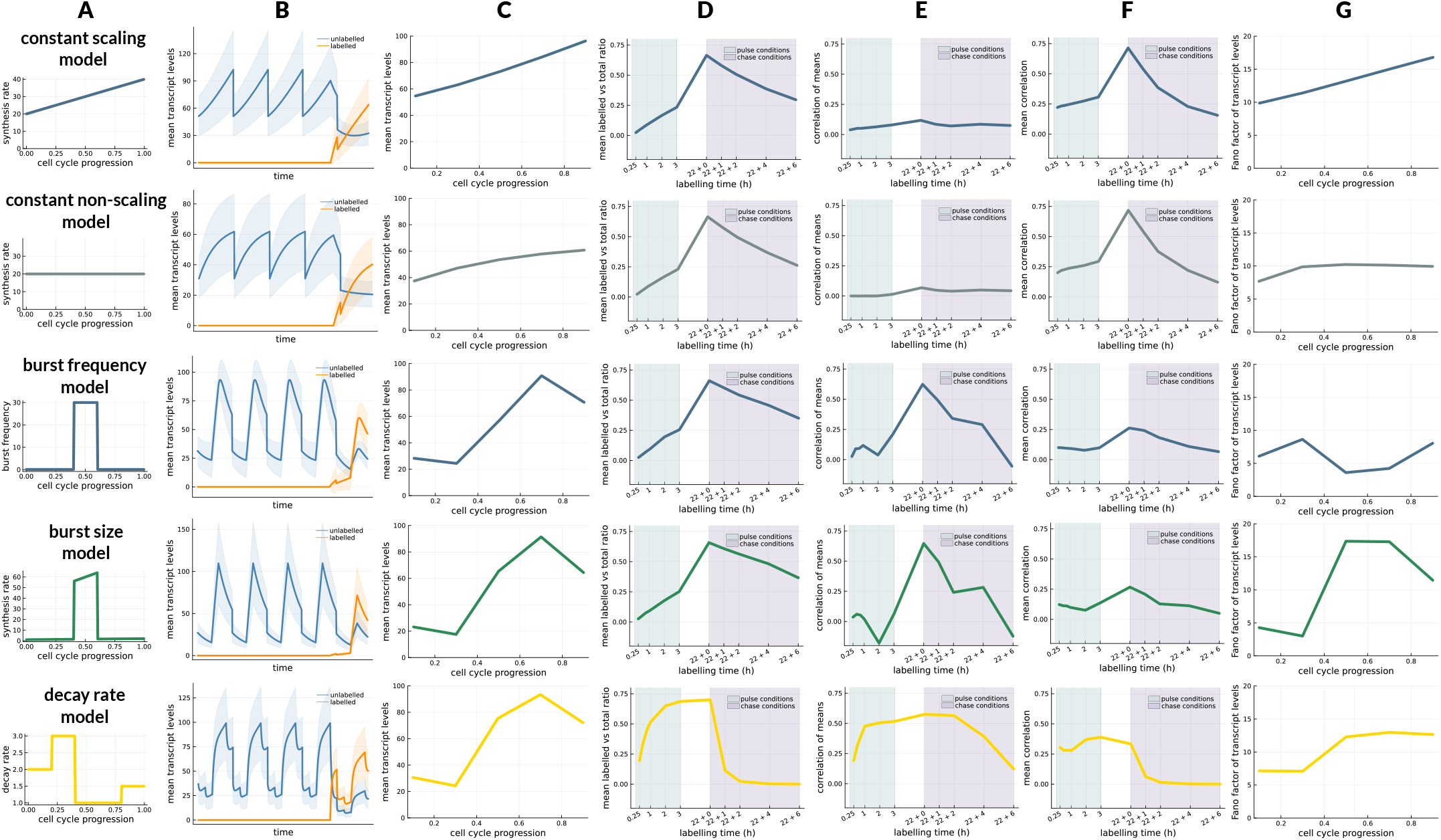
Comparison of summary statistics arising from the different models. (A) Labels of the different models and associated regulation of kinetics (see also Figure 2 (A)). (B) Model realisations for each parameter setup: Mean unlabelled and labelled transcript levels across a single-cell lineage over time, where the shaded region corresponds to the standard deviation of transcript levels. The 1st realisation (row 1) assumes that all kinetic rates are constant along the cell cycle, while the synthesis rate scales linearly with cell size. The 2nd realisation (row 2) assumes identical parameters with the 1st realisation, with the only difference being that the synthesis rate does not depend on cell size. The next 3 realisations (rows 3-5) assume time-varying burst frequency, synthesis rate or decay rate and scaling of synthesis rate with cell size, while the rest of the kinetic rates are constant. The chosen parameter sets in rows 3-5 are such that we obtain almost identical patterns of cell cycle-dependent mean expression (column (C)). (C-G) Summary statistics generated by each model using the cell cycle-dependent and labelling time-dependent moments of unlabelled and labelled mRNA (see Supplementary Methods 1.4 and Tables 1 and 2). Columns (C-E) show statistics that can be computed from mean measurements and are defined in rows (1),(3) and (5) of Table 1. Columns (F-G) show statistics that can be computed from noise measurements and are defined in rows (2) and (4) of Table 1. The constant scaling and non-scaling models (rows 1-2) can be distinguished by the cell cycle-dependent mean and noise statistics (columns (C) and (G)). The varying burst frequency and burst size models (rows 3-4) yield almost identical statistics, but they have a striking difference in the cell cycle-dependent Fano factor. The varying decay rate model (row 5) yields substantially different statistics compared to the other two non-constant models. Distinguishing between constant and non-constant models can be based on correlation coefficients: The correlation of cell cycle-dependent means (column (E)), that measures noise due to cell cycle variation, is higher in non-constant models, while the mean (cell cycle-averaged) correlation, that measures upstream noise due to the bursty gene promoter, is higher in constant models.

**Supplementary Figure S3:**
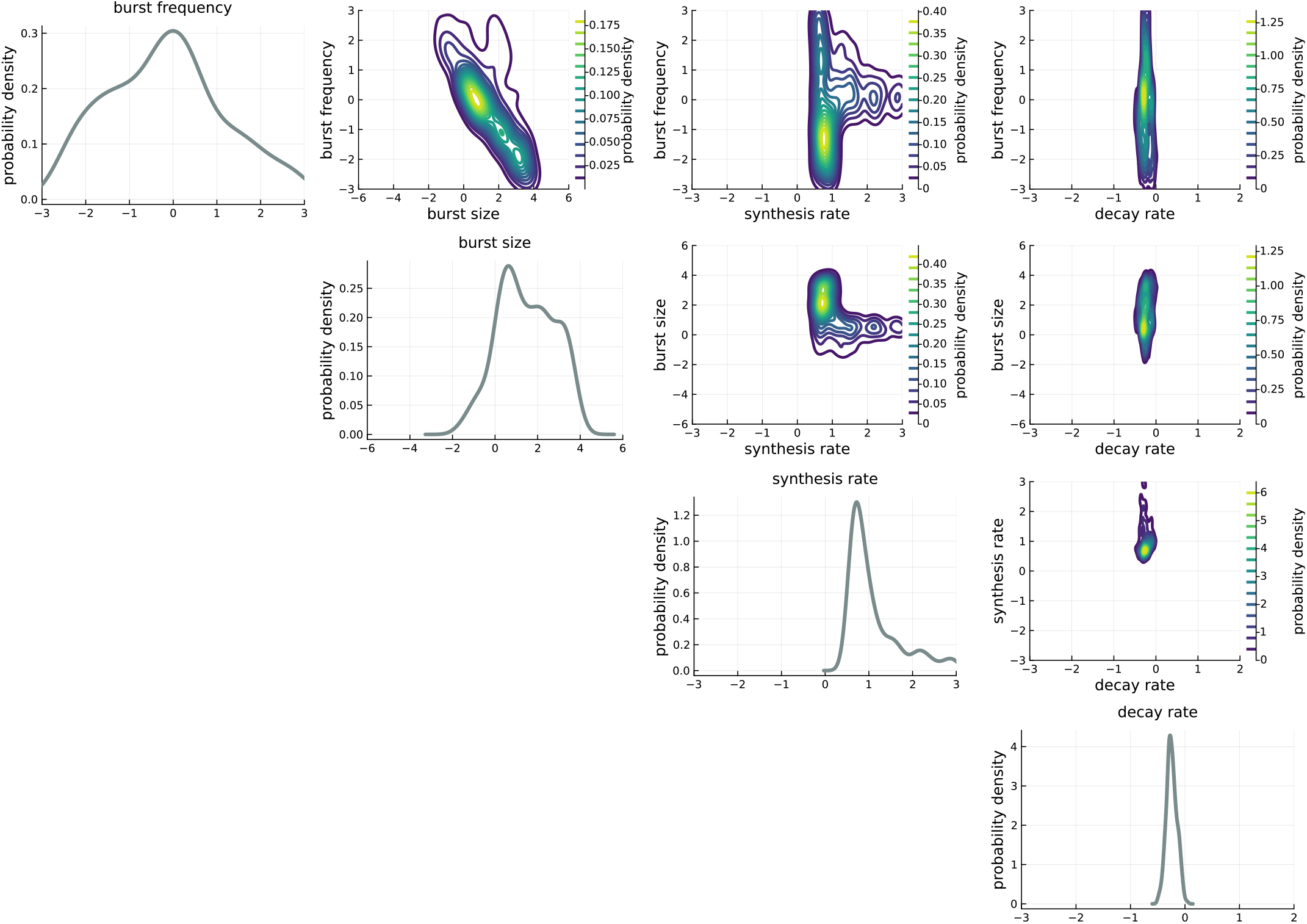
Approximate posterior distributions of parameters for an example gene *SGK1*. The diagonal plots show kernel density estimates of the posterior distributions of burst frequency *k*_on_, burst size *α/k*_off_, synthesis rate *α* and decay rate *γ* in log_10_ scale. The off-diagonal plots show kernel density estimates of the joint posterior distributions of pairs of parameters. All parameters are measured in *h*^−1^, apart from burst size which is unitless.

**Supplementary Figure S4:**
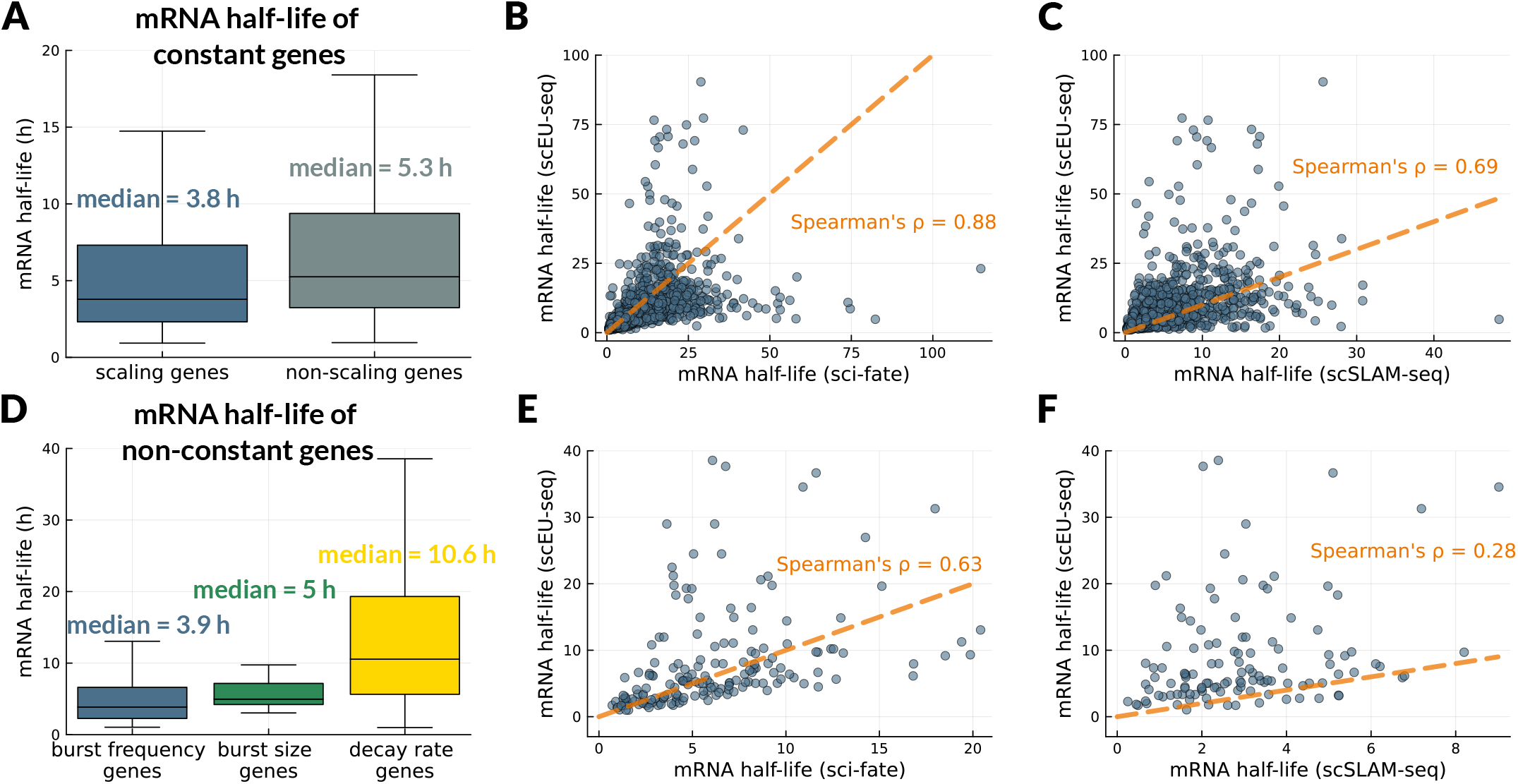
mRNA half-life distributions and comparison with previously reported estimates. (A) Estimated mRNA half-lives of constant scaling and non-scaling genes (in hours), as defined in (10) using the point estimates of decay rate. (B-C) Comparison of half-lives of constant genes with n = 2209 common genes in A549 cells as measured in the sci-fate study [52] and with n = 1565 common genes in NIH-3T3 cells as measured in the scSLAM-seq study [49]. These comparisons exclude half-lives that are larger than 100 hours (less than 2% of constant genes). The orange dashed lines are identity lines. (D) Estimated mRNA half-lives of non-constant genes. To compute the half-lives of varying decay rate genes, we use the average decay rate over the cell cycle. (E-F) Comparison of half-lives of non-constant genes with n = 177 common genes in A549 cells as measured in the sci-fate study and with n = 119 common genes in NIH-3T3 cells as measured in the scSLAM-seq study. These comparisons exclude half-lives that are larger than 40 hours (1% of non-constant genes). The orange dashed lines are identity lines.

**Supplementary Figure S5:**
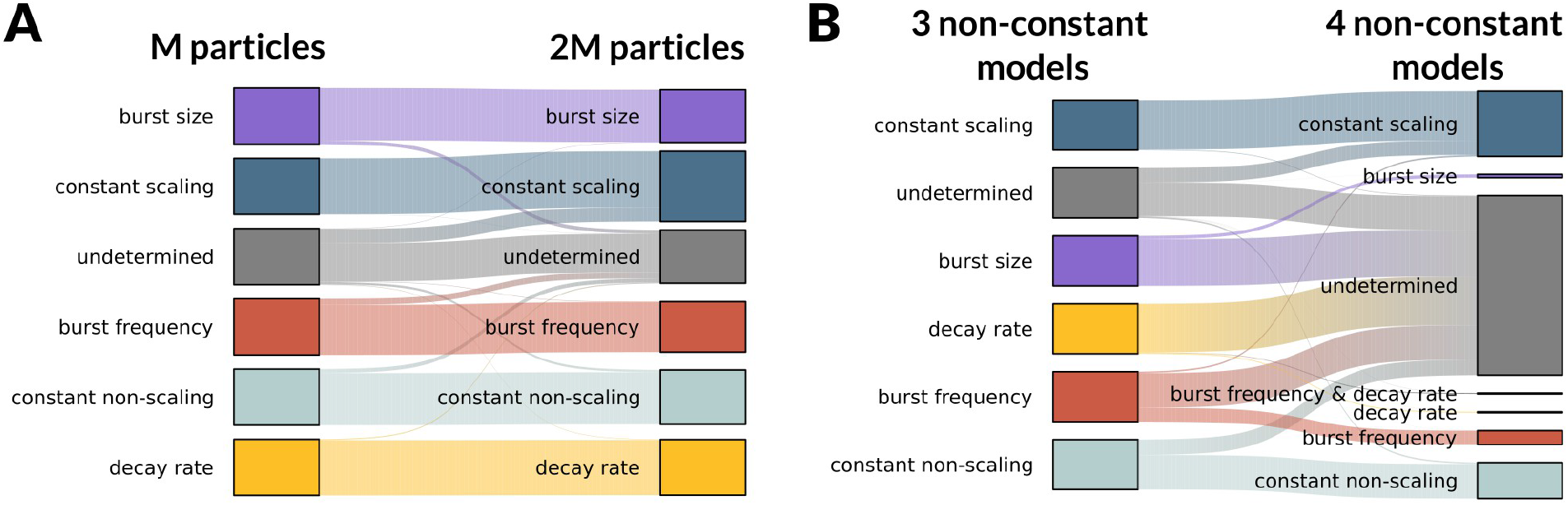
Impact of additional particles and more complex models on model selection. (A) 2-fold increase in the number of particles - simulations (*M* = 5 * 10^6^) does not influence model selection and gene classification. (B) Including an additional model of regulation (varying burst frequency & decay rate) increases uncertainty in selecting between non-constant models.

**Supplementary Figure S6:**
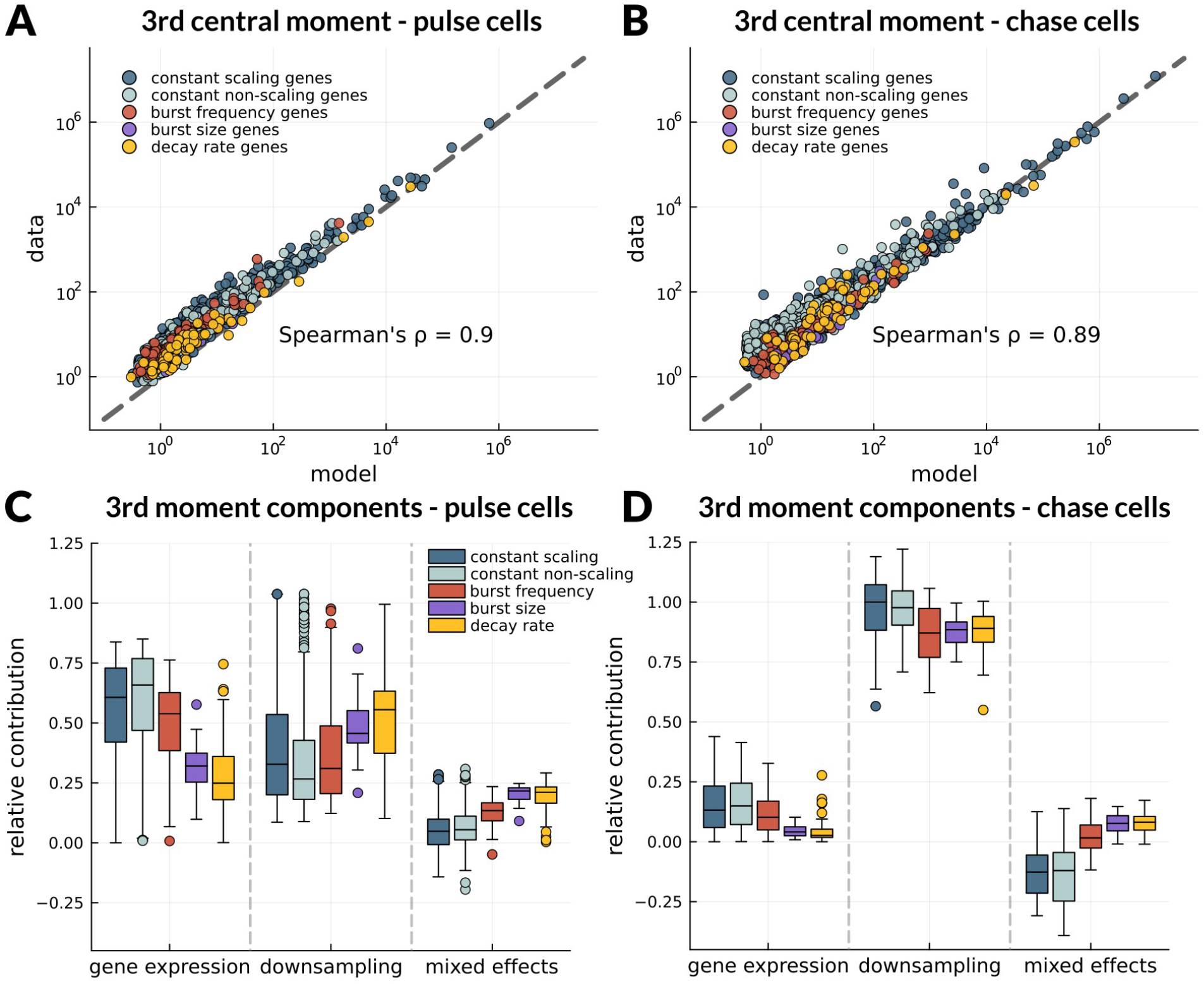
Predictive capacity of the model and sufficiency of summary statistics. (A-B) 3rd order central moments of expression are accurately predicted for pulse-treated and chase-treated cells by the model using gene-specific MAP estimates. (C-D) Relative contributions of biological and technical components to the 3rd central moment for pulse-treated and chase-treated cells, computed using equation (40) in Supplementary Methods. The “mixed-effects” component of the 3rd moment is a covariance that can take negative values.

**Supplementary Figure S7:**
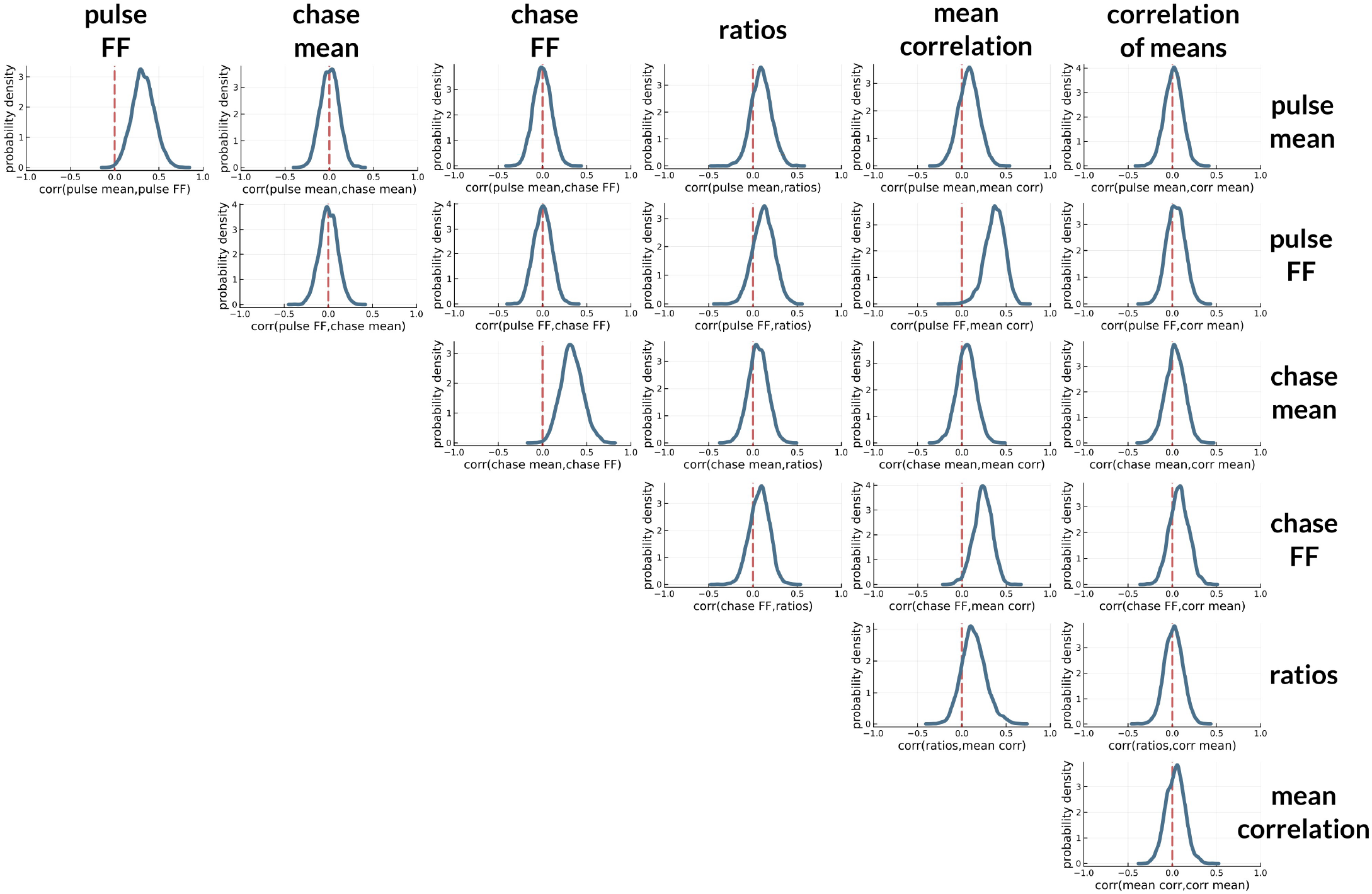
Independence of summary statistics. For each gene, we boot-strapped the data 100 times, we computed the summary statistics across all bootstraps and computed correlations between all different pairs of summary statistics. Inspecting the distributions of these correlations across genes, we find that the majority are centred around 0 or close to 0. Although the small positive correlations can be a result of sample size bias, this result overall suggests that our summary statistics are approximately independent.

**Supplementary Figure S8:**
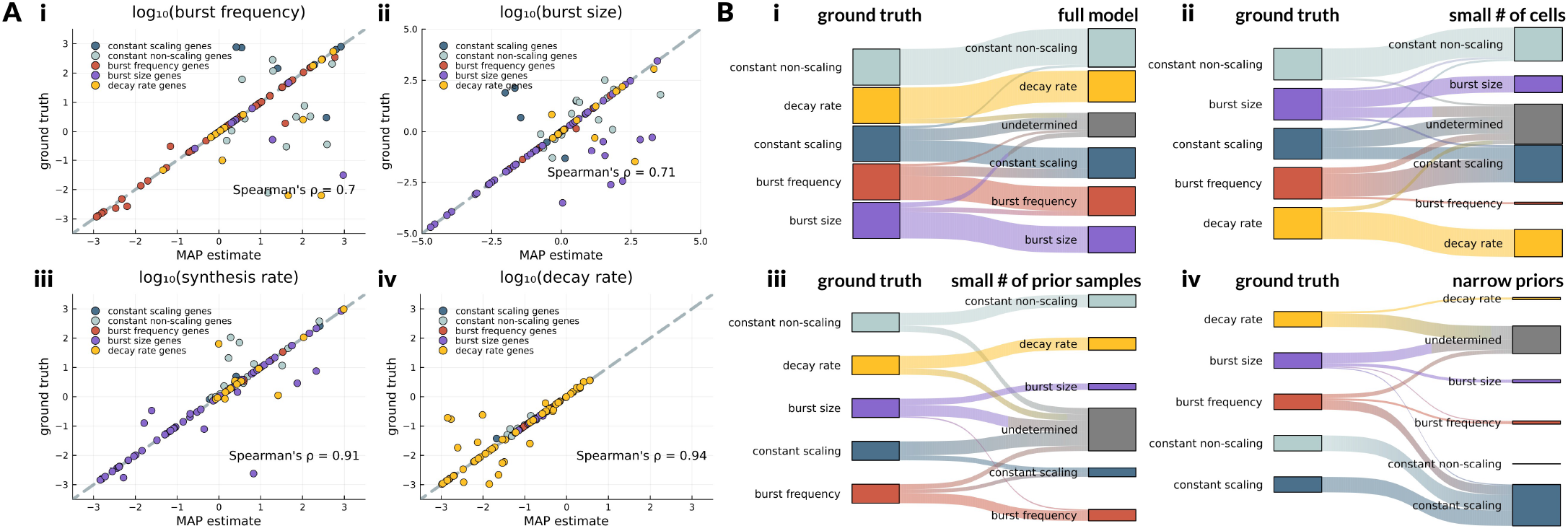
Inference on synthetic data and model robustness. We used Gillespie simulations with ground truth parameters that we sampled from posteriors inferred from the real data to generate datasets of unlabelled and labelled mRNA counts with 2463 cells and 75 genes (Supplementary Methods 1.6). (A) Ground truth values of (i) burst frequency, (ii) burst size, (iii) synthesis rate and (iv) decay rate, are predicted with high accuracy by MAP estimates of the parameters, across all 5 model classes. We plot all parameters corresponding to genes that were classified correctly to the ground truth model class. (B) The robustness in model selection was tested in 4 different scenarios: (i) The full modelling set up (full model) recovers the ground truth model classification with high accuracy. (ii-iii) In the limits of small sample size (small number of cells) and limited parameter sampling (small number of prior samples), the majority of ground truth model classes are recovered successfully, while for a fraction of genes there is significant uncertainty in model selection. Burst frequency genes tend to be classified as constant scaling, which is reasonable as the two models are a special case of each other and can yield similar dynamics. (iv) In the case of narrow priors we observe higher uncertainty in model selection, and many genes are classified as constant scaling, indicating that simpler models are favoured. These results suggest that using a sufficiently wide prior support is essential for accurate model selection.

**Supplementary Figure S9:**
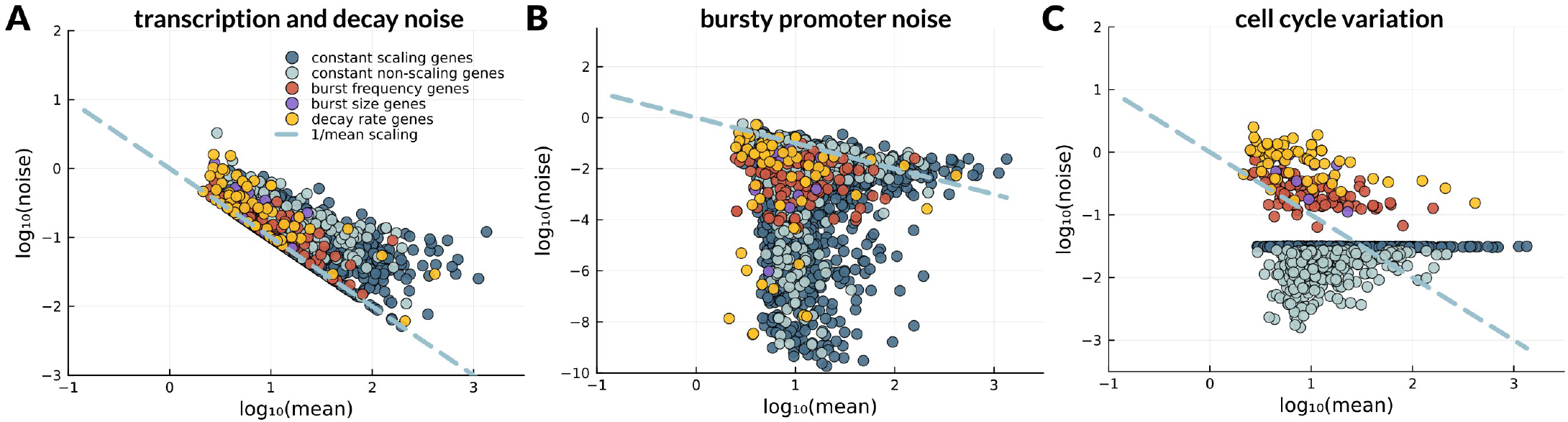
Absolute levels of biological noise sources. Absolute levels of noise dependence on the mean across the different gene classes, for each noise component: (A) transcription and decay noise, (B) bursty promoter noise and (C) cell cycle variation.

**Supplementary Figure S10:**
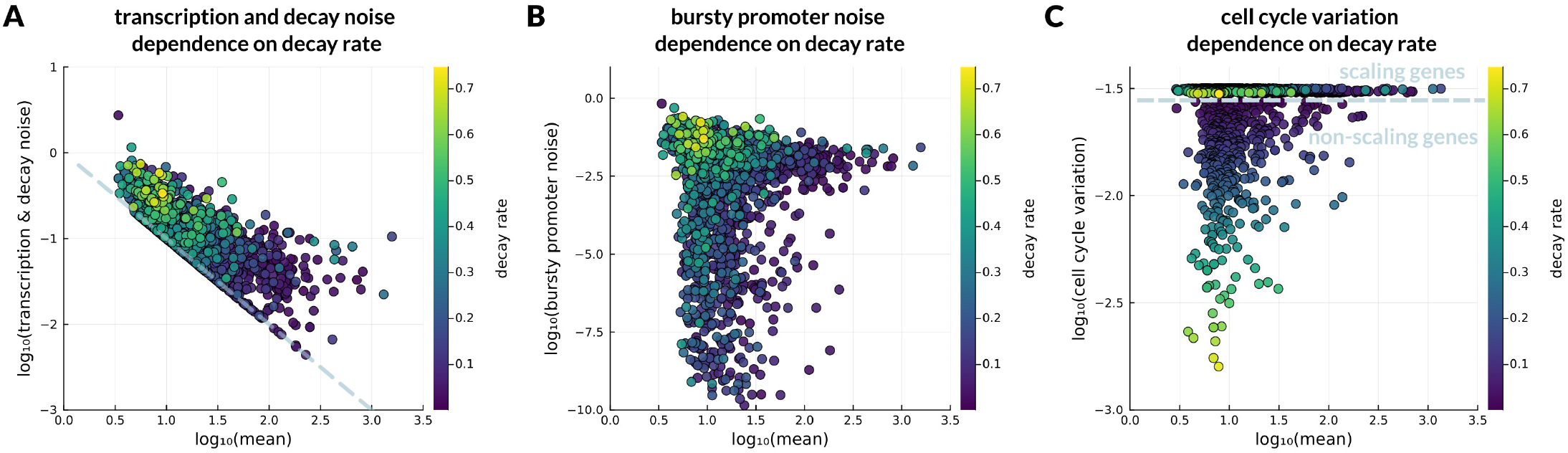
Dependence of noise decomposition on decay rate. We only show decay rates of constant (scaling and non-scaling) genes. Decay rate is measured in *h*^−1^.

**Supplementary Figure S11:**
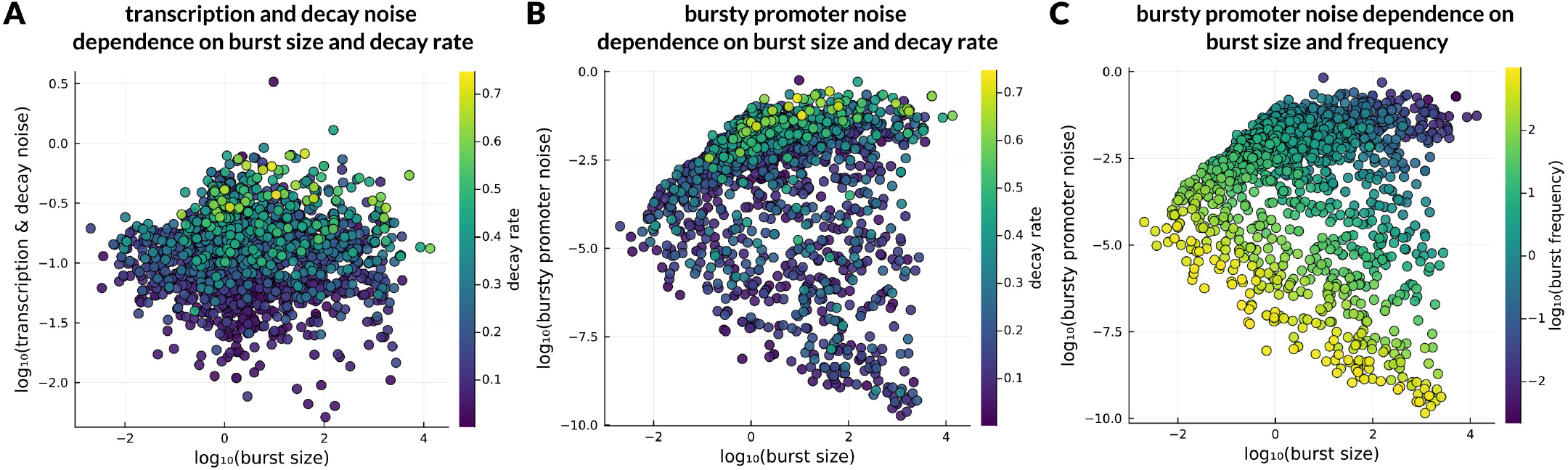
Dependence of noise decomposition on burst size and frequency. Here we only plot kinetic rates associated to constant (scaling and non-scaling) genes. Burst size and frequency are shown in log_10_ scale and decay rate is shown in linear scale. (A) Transcription and decay noise increases with decay rate, while there is no direct dependence on burst size. (B) Bursty promoter noise increases with burst size at the limit of high decay rate. There is no clear dependence observed in the low decay rate regime. (C) Bursty promoter noise decreases with burst size when burst frequency is fixed.

**Supplementary Figure S12:**
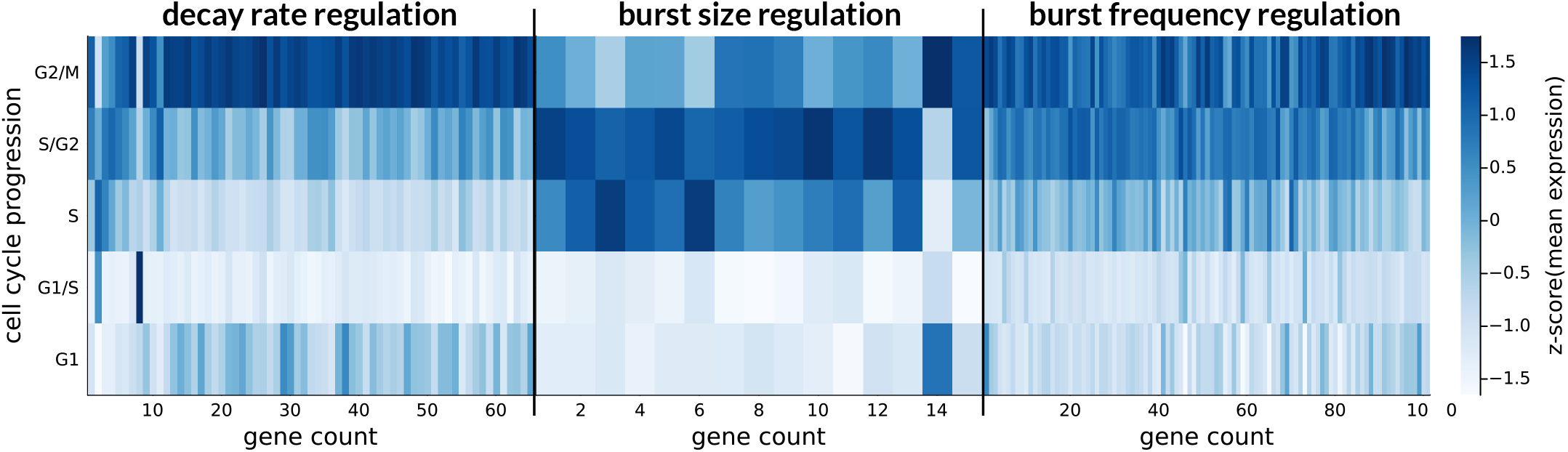
Mean expression profiles of cell cycle-dependent genes. Heatmaps of mean expression z-scores of decay rate genes, burst size genes and burst frequency genes. The ordering of the genes corresponds exactly to the ordering of kinetic rates in the heatmap of Figure 5 (B).

